# Serpins A1/A3 within tumor-derived extracellular vesicles support pro-tumoral bias of neutrophils in cancer

**DOI:** 10.1101/2025.03.06.641593

**Authors:** Maksim Domnich, Ekaterina Pylaeva, Elena Siakaeva, Nastassia Kabankova, Agnieszka Bedzinska, Damian Sojka, Aneta Zebrowska, Marta Gawin, Maren Soldierer, Malwina Rist, Daniel Fochtman, Irem Ozel, Bernd Giebel, Iris Helfrich, Ilona Thiel, Basant Kumar Thakur, Cornelius H.L. Kürten, Helmut Hanenberg, Stephan Lang, Sonja Ludwig, Monika Pietrowska, Jadwiga Jablonska

## Abstract

Neutrophils are known to play an important regulatory role during tumor progression in several types of cancer. However, the mechanisms responsible for their tumorigenic bias and extended lifespan in cancer are not clear to date. This study uncovers a previously unknown mechanism by which tumor-derived small extracellular vesicles (sEVs), via their serpin cargo, reprogram neutrophils to adopt a tumor-supporting phenotype. We demonstrated here an elevated content of plasma sEVs during head and neck cancer progression, and their significant cargo enrichment with inhibitors of neutrophil serine proteases: serpins A1 and A3. Mechanistically, neutrophils educated with serpin-rich tumor-derived sEVs displayed typical pro-tumoral characteristics, including prolonged lifespan and activated CD62L^low^ CD11b^high^ PDL1^high^ phenotype. Functionally, such neutrophils demonstrated a strong ability to promote the epithelial-to-mesenchymal transition of tumor cells. Moreover, such neutrophils induced remarkable suppression of cytotoxic CD8 T cells, significantly reducing their tumor cell-killing capacity. Importantly, serpin cargo was essential for this activity, as serpin-depleted sEVs failed to reprogram neutrophils. These findings again highlight the clinical significance of sEVs and suggest their serpin content as important mediators of pro-tumoral functionality. Targeting the biogenesis or uptake of such immunosuppressive sEVs, or modifying their cargo, could potentially serve as a potent adjuvant anti-cancer therapy.

## Introduction

Neutrophils play an essential regulatory role during cancer progression and their activity is strongly modulated by tumor-released factors, such as cytokines or growth factors^1,2^. We and others observed significant alteration of neutrophil phenotype in tumor-bearing hosts, such as induction of survival^3^, NET formation^4^ and pro-angiogenic activity^5^, but also decreased apoptosis^4^ and significant immunoregulatory capacity^6^. Moreover, neutrophil numbers in the circulation and periphery are significantly elevated in cancer patients, which negatively correlates with patient survival^6–9^. Recently, we demonstrated that while at the initial phase of tumor development, neutrophils exhibit anti-tumoral phenotype, at later tumor stages, these cells become hijacked to support tumor growth^6^. However, the exact mechanism driving this functional switch remain poorly understood. Tumor-released factors (such as chemokines pro-inflammatory and angiogenic cytokines) likely play a critical role in this process^2,10^, with small extracellular vesicles (sEVs) emerging as potential carriers and mediators of communication between tumor cells and the immune system.

Extracellular vesicles are small membrane-bound structures released by cells into their external environment that bear the same properties as the producing cell. These vesicles play crucial roles in intercellular communication by transporting various cargo and are involved in cell-to-cell signaling^11,12^.

Extracellular vesicles have been found to impact immune cell function in various ways: modifying dendritic cell function^13,14^, suppressing T cell proliferation^15,16^ or activating natural killer cells^17^.

In this work, we evaluate the role of cancer-released sEVs in the modulation of neutrophil activity to define the mechanisms responsible for the neutrophil phenotype switch during cancer progression.

## Materials and methods

### Patients

Peripheral venous blood specimens were collected from patients with HNC (45 individuals) treated at University Hospital Essen and healthy volunteers (10 individuals) from 2017 to 2019. The clinical characteristics of the enrolled participants are presented in **Figure 1 A-C**. The patients were scheduled for surgical tumor resection and consented to blood collection for research purposes at the Otorhinolaryngology, Head and Neck Surgery Department of the University Hospital of Duisburg-Essen, Essen, Germany. Patients with other or unknown histological types of tumors like adenocarcinoma, sarcoma, lymphoma, cancer with unknown primary origin (CUP), and tumors associated with specific risk factors like the nasopharyngeal squamous cell carcinoma, which is associated with Epstein Barr Virus, as well as patients that received any cancer-associated therapies prior to the study were omitted. Agreement was received from all individuals. The ethics committee of the University Hospital Essen approved the study.

**Figure 1.**
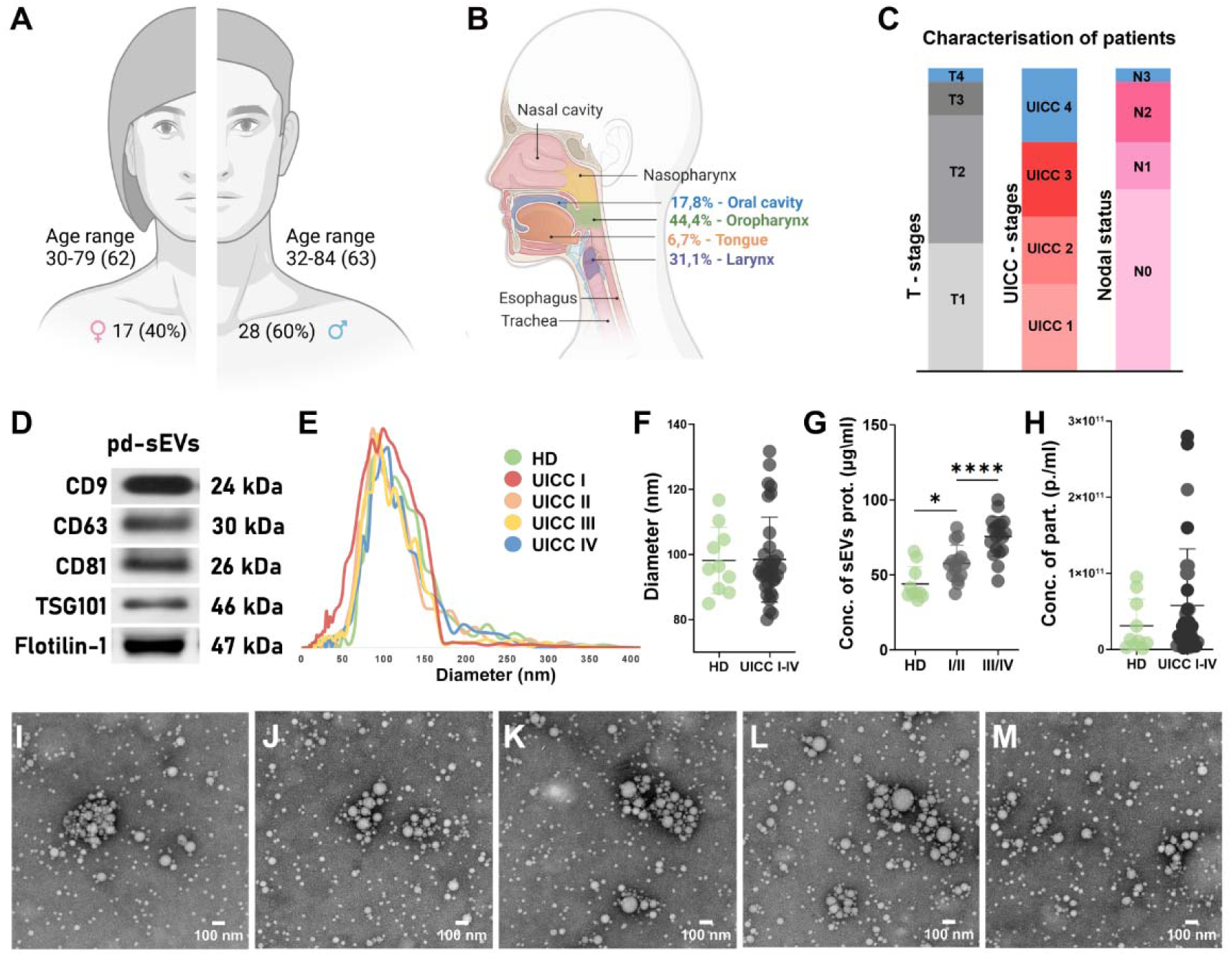
Clinicopathologic profiles of the study participants and characterization of small extracellular vesicles isolated from plasma. Plasma samples were collected from 10 healthy individuals (5 males and 5 females, with a mean age of 38 years) and 45 patients (28 males, 60% and 17 females, 40%, with a mean age of 63 years) (**A**). The patients were classified by the eighth edition of the Union for International Cancer Control (UICC) guidelines according to UICC stage I-IV head and neck squamous cell carcinoma. Primary tumor sites were oropharynx (44.4%) oral cavity (17.8%) including tongue (6.7%) and larynx (31.1%) (**B**). 13 (28.8%) patients were UICC I, 10 (22.2%) UICC II, 11 (24.5%) UICC III and 11 (24.5%) UICC IV (**C**). Using western blotting, the presence of common sEVs markers such as CD9, CD81, CD63, TSG101, and Flotilin-1 was determined in plasma-derived pd-sEVs from the plasma samples of study participants (**D**). Nanoparticle tracking analysis (NTA) of sEVs enriched from plasma samples of HD and patients with low (UICC I/II) and high advanced (UICC III/IV) of HNC reveal similar mean sizes (∼110 nm) (**E, G**) and concentrations (**H**) of EVs. In comparison to cancer patients, healthy donors had significantly lower levels of total protein concentration (HD = 44 µg/ml, UICC I/II = 57.3 µg/ml, UICC III/IV 75.7 µg/ml (**F**). The representative TEM micrographs (I – HD, **J** – UICC **I, K** – UICC II, **L** – UICC III, **M** – UICC IV) of sEVs show intact vesicles ranging in size from 50 to 200 nm, which match the data obtained by NTA. Scale bar 100 nm. For the comparison of two independent groups, the Mann-Whitney *U* test was used. Data are shown as individual values with a median. * p < 0.05, **** p < 0.0001.

### Extracellular vesicles Isolation from human plasma

Peripheral blood was drawn into sodium citrate anticoagulant S-Monovette® (Sarstedt, cat# 02.1067.001), centrifuged at 1,500 x g at 20°C for 10 minutes, plasma was collected and aliquoted in 1-mL tubes, and stored at −80°C until usage. Either using fresh plasma or after thawing of stored plasma, samples were differentially centrifuged 2,000 x g for 10 min at 20°C and then centrifuged 14,000 x g for 30 min at 4°C for pre-cleaning from large microvesicles and apoptotic bodies. Then, plasma was ultrafiltrated using a 0.22-μm filter (Millex, cat# SLGP033RS). Next, the Econo-Pac® chromatography columns (BioRad, cat# 7321010) filled with 10 ml of Sepharose^TM^ CL-2B (Cytiva, cat# 17-0140-01) were prepared. To harvest EVs, 1 ml of prepared plasma was passed through the column. The fourth fraction (F4) was collected for further investigation.

### Cell lines

UT-SCC-50: Human laryngeal squamous cell carcinoma UT-SCC-50 (p53-frameshift, c.920del73/p14 - No transcript/HPV16 E6/E7−) was obtained free of charge from University Hospital Ulm^18,19^.

UM-SCC-10a: Human squamous cell carcinoma of the larynx UM-SCC-10a (p53 - missense, p.245Gly->Cys/p14 - frameshift, c.388delC/HPV16 E6/E7−) was kindly provided by professor Thomas E. Carey (Ph.D., Director, Laboratory of Head and Neck Cancer Biology, Department of Otolaryngology/Head and Neck Surgery University of Michigan, Ann Arbor, USA)^20,21^.

For the Serpin A1 and Serpin A3 CRISPR/Cas9 knockout constructs, DNA oligonucleotides for the gRNAs (CTATGATCTGAAGAGCGTCC for Serpin A1; CGGATTAGCCTCCGCCAACG for Serpin A3, as well as a control gRNA targeting chicken beta-actin were cloned into the lentiviral CRISPR/Cas9 vector LentiCRISPRv2 (Addgene, cat# 52961) using BsmBI restriction enzyme sites as described^22^. VSV-G-pseudotyped replication-deficient infectious lentiviral particles were produced in HEK293T cells as previously described^23^. After filtering through 0.45 μm, the lentiviral particles were used freshly for the transduction of UM-SCC10a cells. After 24 h, the viral supernatant was replaced with a fresh cell culture medium. Three days after transduction, cells were selected with 1µg/ml puromycin for five days and then used for functional studies.

### Cell lines supernatants

The cells were cultivated in DMEM (Gibco, cat# 41966-029) supplemented with 10% of exosome-depleted (ultracentrifuged at 100,000 x g for 18 h) Fetal Bovine Serum (PAN Biotech, cat# P30-3031), 1% of penicillin-streptomycin (Gibco, cat# 15140122), and 100 μM non-essential amino acids (Gibco, cat# 11140050). Cells were grown in a monolayer at 37°C in a humidified incubator with 5% CO_2_. During cultivation, the cell lines were regularly tested for mycoplasma contamination, and the results were negative.

For large production of supernatant enriched by sEVs from the cell lines described above, the 4-layer Nunc™ EasyFill™ Cell Factory™ System (Thermo Fischer, cat# 140004) was used. Cells were split after 72 h of incubation and re-seeded using identical cell numbers. Supernatants were harvested before splitting after 72 h of incubation with confluency around 70-90%.

### Extracellular vesicles Isolation from cell line supernatants

A 50 mL aliquot of cell culture supernatant was centrifuged at 20°C for 10 min at 2,000 × g to separate cells and debris. Then supernatants were transferred to new tubes and centrifuged again at 4°C for 30 minutes at 10,000 x g. Afterward, the supernatants were collected and filtrated using a 50 mL syringe and a 0.22 μm PES bacterial filter (Millipore, cat# SLGP033RS). In the next step, 50 ml supernatant aliquots were concentrated 50 times to 1 ml using Vivacell 100 concentrators (Sartorius, cat# VS2042) at 3,000 x g with a cut-off of 100 kDa. Small extracellular vesicles were isolated using the size exclusion columns in the same way described for human plasma.

### Nanoparticle tracking analysis

Multiple–-Laser ZetaView® f-NTA Nanoparticle Tracking Analyzer (Particle Metrix, Inning am Ammersee, Germany) was used to evaluate the concentration and size distribution of extracellular vesicles. A pre-testing series (140-200 particles/frame) was performed to assess the proper dilution of sEVs fraction 4. The software settings for sEVs measurement have been selected according to the manufacturer’s recommendation. Three repetitions were performed for each sample by scanning 11 cell positions each and capturing 60 frames per position with the following settings: video setting: high; Focus: autofocus; Camera sensitivity for all samples: 92.0; Shutter: 70; Scattering Intensity: 4.0; Cell temperature: 25°C. Captured videos were analyzed by the in-build ZetaView Software with set parameters: Maximum particle size: 1000; Minimum particle size 5; Minimum particle brightness: 20; Hardware: embedded laser: 40 mW at 488 nm; camera: CMOS.

### Transmission electron microscopy (TEM)

To determine the morphology of sEVs, samples were fixed by 1% osmium tetroxide (OsO4) solution (Sigma, cat# 75632-5ML) in proportion 1:1 for 30 min. After that on top of fixated samples (10 μL) were placed formvar-coated TEM grids (EMS, cat# EFCF200-Cu-TC) for 30 min. In the next step, grids were cleaned out of the fixator by transferring to drops of distilled water (10 μL) for 5 min with 3 repetitions. Then, grids were placed into drops of 1% uranyl-acetate (EMS, cat# 100504-336) in 50 % alcohol (10 μL) for contrast staining for 15 min. In the last step samples were washed by transferring to drops of distillated water for 5 min with 3 repetitions. Afterwards, the last drops of water were airdried and ready for TEM. Samples were analyzed under JEM 1400 Plus transmission electron microscope (JEOL, Peabody, MA USA) integrated with TVIPS TemCam-F416 high-sensitive CMOS camera (TVIPS, Martinsried, Germany) allows to take 16Bit digital micrographs at a resolution of 4,096 x 4,096 pixels using the EM-Menu 4.0 extended software. The contrast, brightness, and sharpness of electron micrographs were edited using Adobe Photoshop CC (Adobe, San Jose, CA, USA).

### Western blotting

Ten µL of SEC fraction - F4 were mixed with loading buffer (BioRad) and optionally 100 mM DTT (in the case when reduction conditions were required), then denatured for 5 min at 95°C. PBS was used as a negative control. A mixture of SEC fractions F5 and F6 derived from healthy volunteers was used as a positive control. Samples separation was performed using 12% SDS-polyacrylamide gel electrophoresis followed by wet electro-transfer onto nitrocellulose membranes (Thermo Fisher, cat# 88018). Membranes were blocked for 1 h in 5% non-fatty milk and 0.1% Tween in PBS (for anti-CD9, anti-CD63, anti-TSG101) or in 5% BSA and 0.1% Tween in PBS (for anti-CD81). Primary antibodies were added for overnight incubation at 4°C (anti-CD63: Invitrogen, cat# 10628D, 1:1500; anti-CD9: Santa Cruz Biotechnology, sc-13118, 1:500; anti-CD81: Biorbyt, orb388959, 1:500; anti-TSG101: Becton Dickinson, 612697, 1:800). Analysis of CD63 and CD81 on SDS-PAGE were performed under non-reducing conditions. The membranes were washed five times with Tris-buffered saline with Tween buffer (20 mM Tris, pH 7.5, 150 mM NaCl, 0.1% Tween 20) at 20°C, and then the secondary antibody conjugated with HRP, diluted in the same solution as the primary, was added for 1 h incubation at 20°C. After five washes, WesternBright Sirius HRP substrate (Advansta, cat# K-12043-D10) was used for chemiluminescence detection of the bands according to the manufacturer’s instructions.

### Sample preparation for LC-MS/MS

The samples of sEVs (fraction #4 only) were thawed and centrifuged: 30 min, 16,000 x g, 4°C to remove any suspended matter, then supernatants were collected and subjected to protein precipitation with the use of 10% (m/v) trichloroacetic acid in acetone (precipitant to sample volume ratio of 3:1, incubation at −20°C for 15 h). Next, samples were centrifuged: 30 min, 16,000 x g, 4°C. Supernatants were carefully collected and discarded, and protein pellets were washed twice with a portion of 500 µL of pure ice-cold acetone, each washing followed by centrifugation: 30 min, 16,000 x g, 4°C. Leftovers of acetone were removed via gentle heating of the tubes in a thermoblock (37°C, ca. 10 min). Each protein pellet was subsequently soaked with 20 µL of 0.2% (m/v) RapiGest in 50 mM NH_4_HCO_3_, mixed and boiled at 100°C for 10 min, then cooled down at room temperature and spun down shortly. Next, the samples were subjected to protein quantification with the use of tryptophan fluorescence method as published before^24^. Finally, proteins were reduced with dithiothreitol (final concentration of DTT: 5 mM, heating at 60°C for 30 min), alkylated with iodoacetamide (final concentration of IAA: 15 mM, incubation in darkness for 30 min) and subjected to in-solution digestion using Trypsin/Lys-C Mix (Promega, cat# V5073) with enzyme to protein ratio of 1:50 (m/m). The digestion was carried out at 37°C for 18 h. Next, samples were acidified with trifluoroacetic acid (final concentration of TFA: 0.5%, v/v), and incubated for 45 min at 37°C. This was followed by centrifugation: 13,000 rpm, 20 min, 20°C, and supernatants were carefully collected. Thus, the obtained peptide samples were subsequently purified on C18-StageTips proposed by Rappsilber et al.^25^ and packed with 6 plugs of a C18 extraction disk. Eluates were evaporated to dryness, then reconstituted in water and peptide content was determined with the use of tryptophan fluorescence method; further, samples were acidified with TFA (final acid concentration: 0.1%). Finally, samples from patients belonging to the same cancer progression stage were pooled. Samples derived from healthy donors (HD) were pooled separately. Peptides present in the pooled samples were preconcentrated with the use of Cleanert C18 1 mL SPE cartridges (Agela Technologies, cat# S180501). Eluates were then evaporated to dryness, reconstituted in water and peptide content was determined with the use of tryptophan fluorescence method. Finally, samples were acidified with TFA (final acid concentration: 0.1%) and frozen until LC-MS/MS analysis.

### Protein identification and quantification by LC-MS/MS

Equal amounts of proteins (0.6 μg) from the sEVs samples were uploaded to the Dionex UltiMate 3000 RSLC nanoLC System connected to the Q Exactive Plus Orbitrap mass spectrometer (Thermo Fisher). Next, peptides were separated on a reverse-phase Acclaim PepMap RSLC nanoViper C18 column (75 μm × 25 cm, 2 μm granulation) using acetonitrile gradient (from 4 to 60%, in 0.1% formic acid) at 30°C and a flow rate of 300 nL/min (total run time: 180 min). The spectrometer was operated in data-dependent MS/MS mode with survey scans acquired at the resolution of 70,000 at m/z 200 in MS mode, and 17,500 at m/z 200 in MS2 mode. Spectra were recorded in the scanning range of 300–2000 m/z in the positive ion mode. Higher energy collisional dissociation (HCD) ion fragmentation was performed with normalized collision energies set to 25. A reviewed Swiss-Prot human database was used for protein identification with a precision tolerance of 10 ppm for peptide masses and 0.02 Da for fragment ion masses. A protein was considered positively identified if the search engine found at least two peptides per protein, and if a peptide score reached the significance threshold FDR = 0.01 (as determined by the Percolator algorithm). Furthermore, a protein was classified as ’present’ if detected in at least one sample of a given type. The abundance of identified proteins was estimated using the Precursor Ions Area detector node in Proteome Discoverer. This method calculates protein abundance based on the average intensity of each protein’s three most intense distinct peptides, normalized to the total ion current (TIC). Plasma proteins such as immunoglobulins, apolipoproteins and albumins were considered as a contaminant and excluded from the further comparison (**Tables S2,S3**). The mass spectrometry proteomics data have been deposited to the ProteomeXchange Consortium via the PRIDE^26^ partner repository with the dataset identifier PXD057119.

### Preparation of EVs-depleted cell culture medium

Cell culture medium was prepared according to the protocol of Thery et al.^27^. Briefly, medium complemented with all required nutrients and 20% (v/v) FBS was ultracentrifuged 18 h at 100,000 × g at 4°C (45Ti rotor, Beckman Coulter, cat# 339160). Then, the collected supernatant was sterilized with the use of a 0.22-μm syringe filter and stored at 4°C. The desired concentration was obtained by diluting the complete medium without FBS.

### Human neutrophil and T cell isolation

Peripheral blood was drawn into 3.8% sodium citrate anticoagulant monovettes (Sarstedt, cat# 02.1067.001) and mixed 1:1 with PBS (Gibco, cat# 14040133) before separation by density gradient centrifugation (Biocoll density 1077 g/mL, Merck, cat# L6115) 30 min at 300 x g (without a break) at 20°C. The mononuclear cell fraction was collected, resuspended in buffer (PBS, 0.5% bovine serum albumin (R&D, cat# 5217), 2 mM EDTA (Sigma-Aldrich, cat# E9884)) and centrifuged at 300 x g 4°C (repeated 3 times to minimize the contamination with platelets). T cells were isolated from the mononuclear cells using the immunomagnetic separation (pan T cell negative selection kit, Miltenyi, cat# 130-096-535) following the manufacturer protocol. Isolated T cells (purity ∼95%) were stored on ice in RPMIc medium (RPMI 1640 Medium (Gibco, cat# 11875093), 10% of exosome-depleted (ultracentrifuged at 100,000 x g for 18 h) Fetal Bovine Serum (PAN Biotech, cat# P30-3031) and 1% penicillin-streptomycin (Gibco, cat# 15140122)) for 1 h. Neutrophils were isolated from the polymorphonuclear cell fraction by sedimentation over 1% polyvinyl alcohol (Sigma-Aldrich, cat# 360627) for 30 min at 20°C, followed by hypotonic lysis of erythrocytes with 0.2% NaCl solution for 1 min at 20°C and reconstitution of osmolarity with 1.2% NaCl. Isolated neutrophils (purity ∼95%) were resuspended in RPMIc medium at 20°C to reach the concentration of 4×10^6^ cells/ml.

### Internalisation of sEVs by neutrophils

The ability of tumor-derived sEVs to be internalized by neutrophils was investigated by examining the uptake of sEVs isolated using SEC and labeled with CFSE dye (Biolegend, cat# 423801). To perform CFSE staining, 15 μl of a 40 μM CFSE solution was added to 100 μL of PBS, containing 10 μg of sEVs. This mixture was then incubated for 45 minutes at +37°C, 5% CO_2_. To eliminate unbound CFSE dye, we employed size exclusion chromatography as described previously. Concurrently freshly isolated neutrophils were co-incubated together with DID (Biotium, cat# 60014-5mg) to label the cell surface and PureBlu Hoechst 33342 (Biorad, cat# 1351304EDU) to visualize the DNA, following the manufacturer’s instructions for both stains. Next, 5×10^5^ of neutrophils were applied to a glass-bottom 96-well plate (MatTek, cat# P96G-1.5-5-F) precoated with poly-D-lysine 1 mg/mL (Sigma-Aldrich, cat# P0899-10MG) for 4 hours at +37°C, 5% CO_2_. The plates were then centrifuged for 5 minutes at 300 x g to ensure cell adhesion. After removing the supernatant, 100 μl of PBS containing 10 μg of sEVs was added to the wells, followed by a 2-hour incubation at +37°C, 5% CO_2_. The plates were washed twice with 200 μl of PBS and fixed with 4% paraformaldehyde (Alfa Aesar, cat# 43368). Images were collected on Zeiss LSM 710 combined with ELYRA PS.1 Super-resolution and TIRF (total internal reflection fluorescence) module using 100X, 1.46 NA Zeiss objective and GaASP detector (Carl Zeiss, Germany). Cell mid-sections were acquired at 12-bit image depth with line averaging (setting 4) and XY pixel size 83 nm in the following channels: DID (Ex: 646 nm; Em: 663 nm), CFSE (Ex: 488 nm; Em: 490–553 nm), PureBlue Hoechst (Ex: 350 nm; Em: 461 nm). Images were scaled to 8-bit RGB identically in Zen software (Carl Zeiss) and exported in JPEG format.

### Education of neutrophils using EVs

To investigate changes in neutrophil phenotype and functions under the influence of tumor-derived sEVs (td-sEVs), neutrophils (5×10^5^ in 200 µl of RPMIc) were co-incubated with 10 µg of sEVs isolated from tumor cell lines in 50 µL of PBS (or PBS alone as a negative control) in 24 well flat-bottom plates for 1h (further co-incubation with tumor cells and T cells, see below) or 18h (for evaluation of survival and phenotypical changes, see below) at 37°C prepared as described above.

### Survival assay

Spontaneous apoptosis of neutrophils isolated from human blood neutrophils was determined by Annexin V/7-aminoactinomycin (7AAD) apoptosis detection kit (BD, cat# 559763) after 18 hours of incubation in RPMIc medium, with or without sEVs. The percentages of alive (annexin V^−^/7-AAD^−^), apoptotic (early apoptosis, annexin V^+^ /7-AAD^−^), and dead by apoptosis (late apoptosis, annexin V^+^/7-AAD^+^) cells from all single cells were estimated.

### Flow cytometry

Single-cell suspensions were stained with eBioscience Fixable Viability Dye (eBioscience, cat# 65-0865-14) and specific antibodies (see supplementary). Staining was performed for 30 min at 4°C. Appropriate isotype control antibodies were used in an additional reaction set. The specific marker expression levels are shown as mean fluorescence intensity (MFI) and calculated by subtraction of corresponding isotype control MFI from specific marker MFI.

### Induction of EMT by tumor-derived sEVs-primed neutrophils

Isolated neutrophils were co-incubated with td-sEVs or PBS as a negative control (as described above) for 1 h. Td-sEVs not internalized by neutrophils were removed by washing using PBS and centrifugation at 300 x g for 5 minutes, 20°C, repeated 3 times. Primed neutrophils were mixed with UT-SCC-50 tumor cells in the ratio 10:1 (150,000 neutrophils to 15,000 tumor cells) in 200 µL of DMEMc in a 48-well plate in duplicate for each biological replicate and incubated for 24 h at 37°C 5% CO_2_. Then the wells were washed gently with PBS to remove neutrophils and dead tumor cells. The UT-SCC-50 cell line was preferred due to its early stage and more prominent epithelial phenotype, which is suitable for investigating epithelial-to-mesenchymal transition (EMT).

To evaluate the morphological changes of the tumor cells driven by td-sEVs-primed neutrophils, well-alive adherent tumor cells remaining in the were fixed with 1 mL ice-cold 96% methanol for 1 h at −20 °C followed by staining with 0.01% water solution of gentianine (Santa Cruz, cat# CAS 439-89-4) for 10 min. The plate was washed with dH_2_O and left to dry overnight. The amount and morphology of tumor cells after co-incubation with td-sEVs-primed neutrophils were evaluated with AMG EVOS digital phase contrast inverted microscope (in 4-8 fields of view, 4 biological replicates) and analyzed by FIJI (ImageJ) software (https://fiji.sc/).

To evaluate the EMT gene reprogramming of tumor cells induced by td-sEVs-primed neutrophils, remaining in the well alive adherent tumor cells were stored in 50 µL of RNA*later* solution (Invitrogen, cat# AM7020) at −20°C. The RNA was isolated using Qia Shredder (Qiagen, cat# 79656) and RNeasy Mini Kit (Qiagen, cat# 74106) and the cDNA was produced using the Superscript II Reverse Transcriptase Kit (Invitrogen, cat# 18064014). qRT-PCR was performed at 60 °C annealing temperature using the following primers; a housekeeping gene, beta-actin ACTB was used (VIM_f AGGCAAAGCAGGAGTCCACTGA, VIM_r ATCTGGCGTTCCAGGGACTCAT, TJP1_f GTCCAGAATCTCGGAAAAGTGCC, TJP1_r CTTTCAGCGCACCATACCAACC, ACTB_f CACCATTGGCAATGAGCGGTTC, ACTB_r AGGTCTTTGCGGATGTCCACGT) The mRNA expression was measured using the Luna Universal qPCR Master Mix (BioLabs, cat# M3003X). Relative gene expressions were calculated by 2^-ΔCt formulations in biological replicates n=4-6.

### Induction of T cell exhaustion by td-sEVs-primed neutrophils

Isolated neutrophils were co-incubated with tds-sEVs or PBS as a negative control (as described above) for 1 h, not internalized by neutrophils td-sEVs were removed with washing using PBS and centrifugation at 300 x g for 5 minutes, 20°C, repeated 3 times. Primed neutrophils were co-incubated with autologous T cells at a ratio of 1:1 (150,000 neutrophils to 150,000 T cells) in 200 µL of RPMIc containing aCD3 (Biolegend, cat# 300402), aCD28 (Biolegend, cat# 302902), and rhIL-2 (Peprotech, cat# 200-02) in 48-well plate and incubated for 24 h at 37°C 5% CO_2._ After priming of T cells by neutrophils, UT-SCC-50 tumor cells were added in proportion 1:10 (15,000 tumor cells in 50 µl of DMEMc) to the wells and further incubated for 24 h at 37°C 5% CO_2._ Then the supernatant and not-adherent neutrophils and T cells were collected, exhaustion state of T cells (alive CD4+ and CD8+) was estimated by the expression of PD1 (Biolegend, cat# 367412) using flow cytometry. The killing capacity of stimulated T cells was assumed to be negatively proportional to the density of the remaining alive adherent UT-SCC-50 tumor cells and was evaluated with AMG EVOS digital phase contrast inverted microscope (in 4-8 fields of view, 4 biological replicates) and analyzed by FIJI (ImageJ) software (version 2.9.0).

### Analysis of the data deposited in the Gene Expression Omnibus databases

We used previously published microarray data deposited in the Gene Expression Omnibus databases GSE65858^28^, GSE83519 (**Table S1**).

### Bioinformatical and statistical analysis

The Perseus software (version 1.6.14) was used for performing statistical and bioinformatical analysis of the mass-spectrometry-based proteomic data. To explore the differences between healthy and patient samples, we used only proteins identified in at least 80 % of samples in each group. Imputation of missing values was made based on the default parameters of the Perseus software. In order to determine the difference in the expression of certain proteins, the LFQ intensity (LFQI) was used as an indicator of the abundance of each protein across the samples. It is calculated based on the intensity with which peptides are detected and on pairwise comparisons of these intensities in any two samples. The resulting LFQI values satisfy the individual protein ratios among samples and preserve the summed-up raw peptide intensities over all samples. Using an ANOVA Multiple-sample Test of the log2 transformed LFQI values, we searched for statistically significant differences in protein abundance. In addition, to estimate the magnitude of such differences, we calculated the fold changes as the ratio of median LFQI values with which a protein was detected in the compared groups. Functional Enrichment analysis was performed based on Cytoscape^29^ (https://cytoscape.org/) open source software platform using EnrichmentMap^30^ (https://apps.cytoscape.org/apps/enrichmentmap) and AutoAnnotate (https://apps.cytoscape.org/apps/autoannotate) applications. Hierarchical cluster analysis was performed to visualize differences in expression patterns using Morpheus software (https://software.broadinstitute.org/morpheus). ShinyGO was used for visualization of overrepresented pathways^31^ (http://bioinformatics.sdstate.edu/go/). Volcano plots were generated using VolcaNoseR^32^ (https://huygens.science.uva.nl/VolcaNoseR2/). Statistical analyses were performed using Kruskal-Wallis ANOVA for multiple comparisons with the Bonferroni correction, and Mann-Whitney U-test for two independent samples, and Whilcoxon test for dependent samples; correlations were analyzed with Spearman R test (visualization via https://www.bioinformatics.com.cn/^33^). The analysis of the Vesiclepedia database was performed using FunRich^34,35^ (version 3.1.4) (http://www.funrich.org/) Kaplan-Meier curves for the survival function were compared via log-rank –test. Data are shown as a median with an interquartile range or as individual values with a median. P<0.05 was considered significant.

## Results

### Elevated content of sEVs in plasma during head and neck cancer progression

To compare amount of sEVs between head and neck cancer (HNC) patients and healthy individuals, we isolated small extracellular vesicles (sEVs) from the plasma of patients at different stages of HNC disease as well as from healthy donors (HD) (**Figure 1 A-C, Table S2**) as described previously^36–38^. The concentration, size, and quality of isolated sEVs were evaluated in agreement with MISEV2023^39^. The presence of the common sEVs markers (CD9, CD63, CD81, TSG101, and Flotilin) was confirmed using Western blotting (exemplified image **Figure 1 D**). To assess if the biophysical properties of such sEVs are altered in HNC, we compared the size of sEVs between HNC and HD individuals using nanoparticle tracking analysis. No significant variations were observed in the average size of sEVs (**Figure 1 E, F**). However, the amount of protein carried in plasma-derived sEVs was significantly increased in advanced stage sEVs (UICC stage III/IV), as compared to sEVs from an early stage cancer (UICC stage I/II) or HD (**Figure 1 G**). In addition, we did not observe significant differences in concentration of particles between groups (**Figure 1 H**). size and purity of isolated sEVs were confirmed by TEM (**Figure 1 I - M**), showing intact vesicles ranging in size from 50 to 200 nm, which matched the data obtained by NTA (**Figure 1 E**).

### sEVs prepared from plasma of HNC patients are enriched in serpins

To identify possible tumor-associated proteins in sEVs, which might influence cancer progression, we investigated changes in the proteome landscape of the plasma-derived sEVs (pd-sEVs) using LC-MS/MS with label-free quantification (LFQ), and compared HNC sEVs to healthy donors. A total of 125 differentially expressed proteins were identified in such sEVs (**Figure S1, Table S3**). We could detect several members of the serpin family, such as SERPINA1, SERPINA3, SERPINC1, SERPIND1, SERPINF2, and SERPING1, which were upregulated in plasma-derived sEVs of HNC patients, especially in the advanced cancer stages (**Figure 2 A-C, Sup. Figure S2**). In agreement, we observed elevated expression of the corresponding genes in HNC tumor tissue compared to healthy mucosa (**Figure 2 D-I**) (data from open assess database GSE83519, **Table S1**).

**Figure 2.**
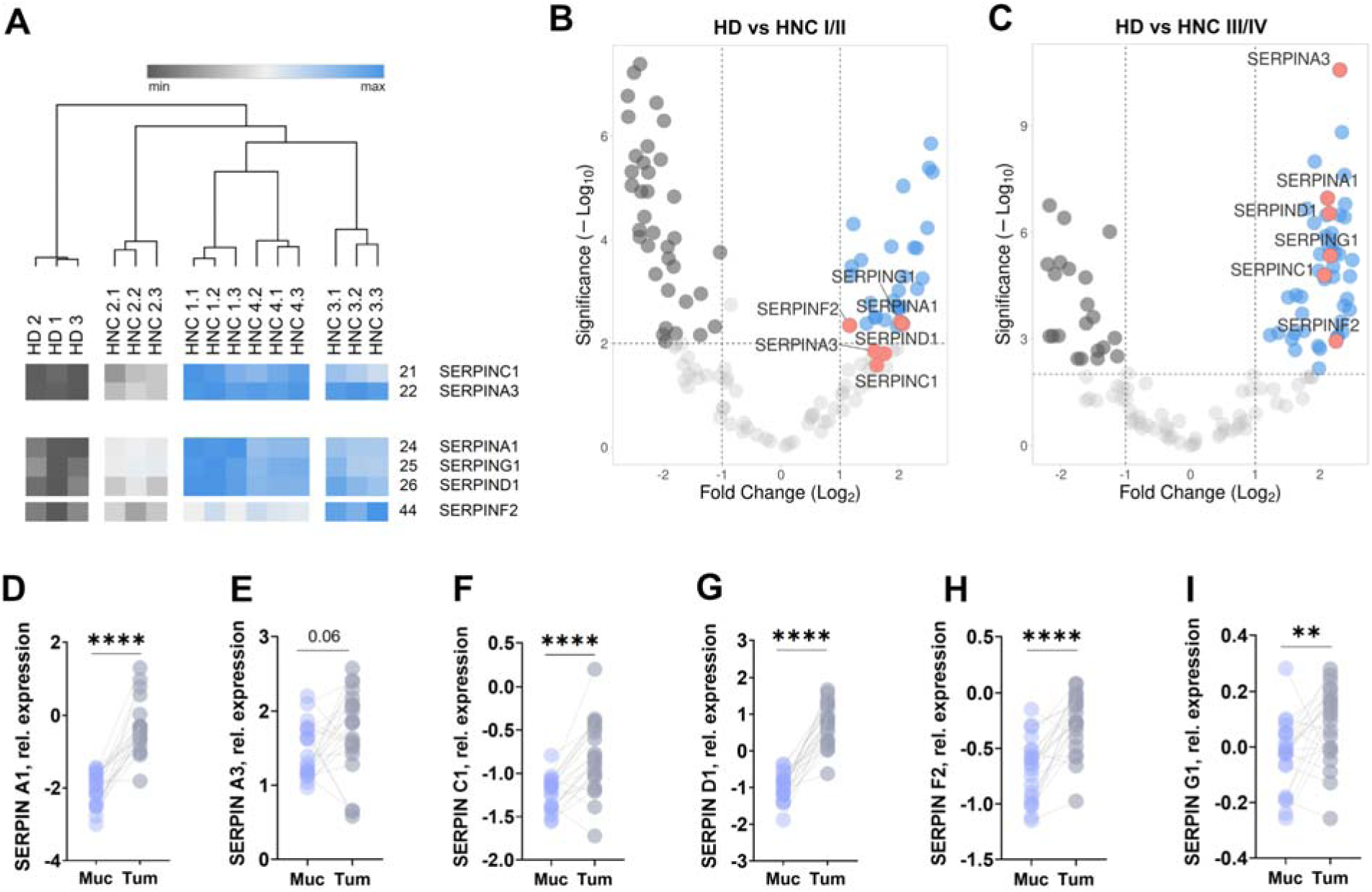
Serpin expression in plasma-derived sEVs and tissues significantly changed during cancer progression. The heatmap (**A**) illustrates the expression intensities of proteins significantly regulated in comparison to healthy controls. Samples (i.e., protein digests) from each group presented on the heatmap (HD 1-3, HNC 1.1-1.3, etc.) consist of protein digests pulled in equal concentrations within groups, and corresponding to the stages of disease progression (HD or UICC I-IV). The relative abundance of each protein was normalized based on a z-score and correlated with the depth of color in the heat map from dark grey (the lowest expression) to dark blue (the highest expression). The dendrogram represents the affinity between the represented groups. Volcano plots (**B, C**) illustrating fold change (log2, x-axis) vs FDR-adjusted P value (Significance, −log10, y-axis) of the plasma-derived sEVs. Data are represented as results obtained for the control group - healthy donors relative to UICC I\II and UICC III\IV. Changes were considered significant at p < 0.01 and fold change > -1.5 (downregulation in grey) or < 1.5 (upregulation in blue) compared with the control group. The gray perpendicular puncture line indicates the border between proteins that have passed a significant value. Analysis of the open-access database GSE83519 shows relative expression levels of various serpin proteins in tumor tissue (Tum) compared to healthy mucosa (Muc) (**D-I**) demonstrated the elevated expression of Serpin A1, A3, C1, D1, F2, and G1 in tumor tissues. For the comparison of two independent groups, the Mann-Whitney U test was used, for the comparison of dependent samples Wilcoxon test was used. Data are shown as individual values with a median. ** p < 0.01, **** p < 0.0001.

### Proteome composition of tumor-derived sEVs confirms enrichment in serpins

Plasma-derived sEVs represent a heterogeneous mixture of sEVs derived from tumor cells, but also other cells present in the circulation, such as immune cells. To confirm the presence of serpins in tumor-derived sEVs (td-sEVs), we purified sEVs derived from HNC cell lines from different tum r stages: UT-SCC-50 (early stage, T2N0) and UM-SCC-10a (advanced stage, T3N0M0). The presence of sEVs markers such as CD9, CD81, TSG101, and Flotilin, but also serpins A1 and A3 was confirmed using Western blotting (**Figure 3 A**). In agreement with results obtained for plasma-derived sEVs, no size (**Figure 3 B, C**) and particle amount (**Figure 3 D**) differences between sEVs from tumor cells at different cancer stages were observed. However, here again a significant increase in the protein cargo of sEVs isolated from the advanced stage cancer (UM-SCC-10a) was detected in comparison to sEVs from an early stage (UT-SCC-50) (**Figure 3 E**).

**Figure 3.**
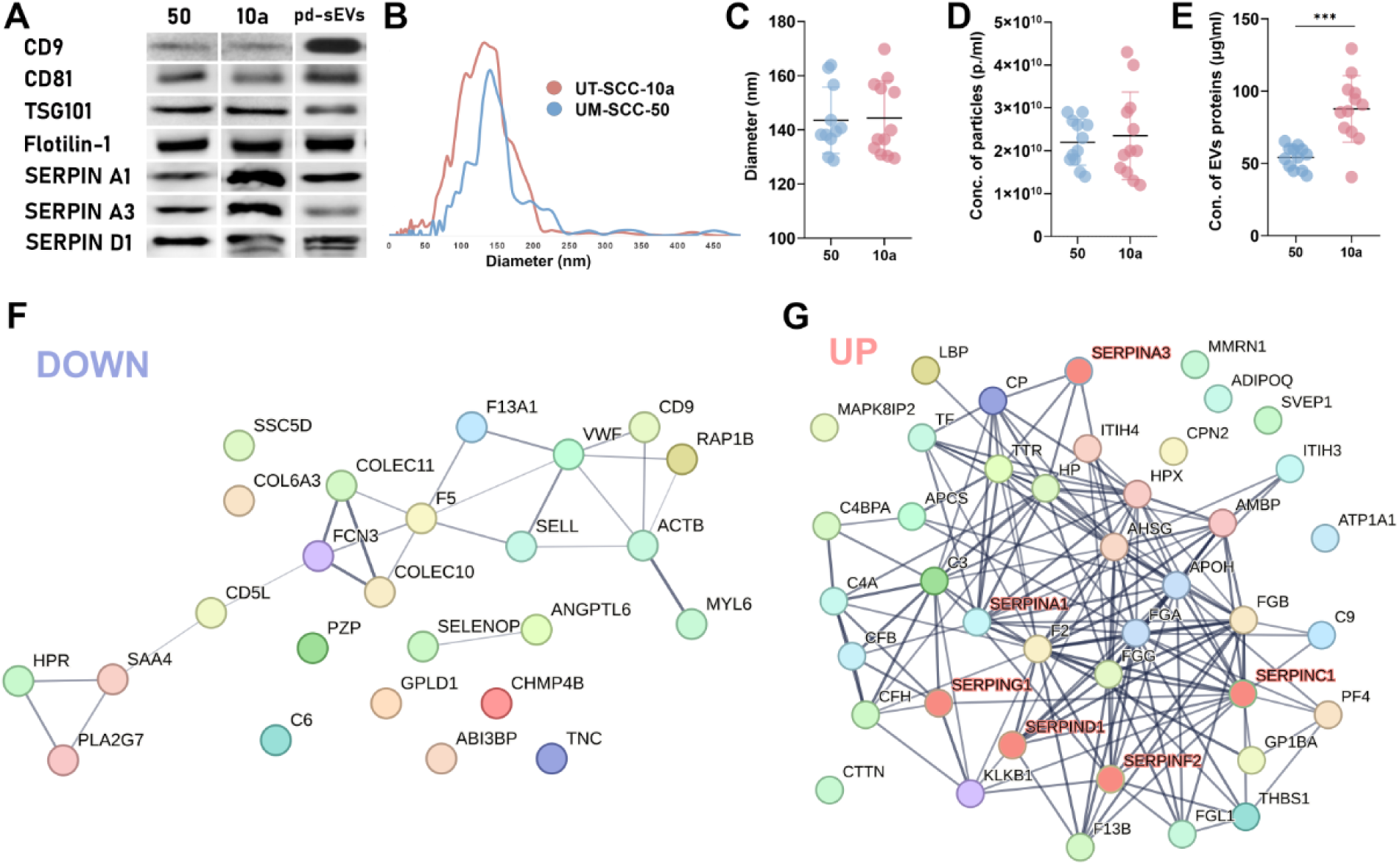
Small Extracellular Vesicles (sEVs) from Early-Stage (UT-SCC-50) and Advanced-Stage (UM-SCC- 10a) HNC cell lines and plasma show enrichment in serpins. Western blotting confirmed the presence of common markers such as CD9, CD81, CD63, TSG101, Flotilin-1 and SERPIN A1, A3 and D1 in sEVs derived from UM-SCC50 (50) and UT-SCC-10a (10) and plasma (pd-sEVs) (**A**). Nanoparticle tracking analysis (NTA) of sEVs isolated from tumor HNC-derived cell lines UT-SCC-10a (pink) and UM-SCC50 (blue) revealed similar mean sizes (**B, C**) and concentrations (**D**). Total protein load of sEVs from advanced cancer stage (UM-SCC-10a) samples was significantly higher in comparison to sEVs from an early stage (UT-SCC-50) (**E**). STRING-based enrichment analysis of proteins that are downregulated (**F**) and upregulated (**G**) in pd-sEVs from HNC patients compared to those from healthy donors. Among others, we could identify several members of the serpin family (SERPINS A1, A3, C1, G1, D1, F2) (**G**), thereby validating their association with the tumor. For the comparison of two independent groups, the Mann-Whitney *U* test was used. Data are shown as individual values. *** p < 0.005.

A detailed analysis of sEVs proteome of patients shows the presence of proteins involved in several signaling pathways potentially involved in the modulation of neutrophil activity, such as metabolic process regulation (influences tumorigenic bias of neutrophils), apoptotic signaling, kinase phosphorylation and neutrophil-mediated immunity (**Sup. Figure. S3**). Inter alia, two main protein clusters were altered: either downregulated or upregulated (including serpins) during cancer progression (**Figure 3 F, G**). Serpins are a family of proteins with inhibitory activity towards neutrophil proteases. Dysregulated pathways involved in neutrophil degranulation play an important role in HNC progression^40^. We focused our attention on serpins A1 and A3, which are both known to inhibit neutrophil granule components elastase and cathepsin G, respectively^41^. Furthermore, the gene signature of neutrophil degranulation (**Sup Figure S4**), among others, is associated with poor prognosis in patients with HNC^40^.

### Tumor-derived sEVs promote survival and pro-tumoral phenotype of neutrophils

To evaluate if td-sEVs could influence neutrophil activity and promote their pro-tumoral phenotype, we educated naïve neutrophils using td-sEVs from HNC cell lines. We observed that neutrophils efficiently internalized td-sEVs (**Figure 4 A -D**), which resulted in significant changes in neutrophil proteome and functionality. (**Figure 4 E - H**).

**Figure 4.**
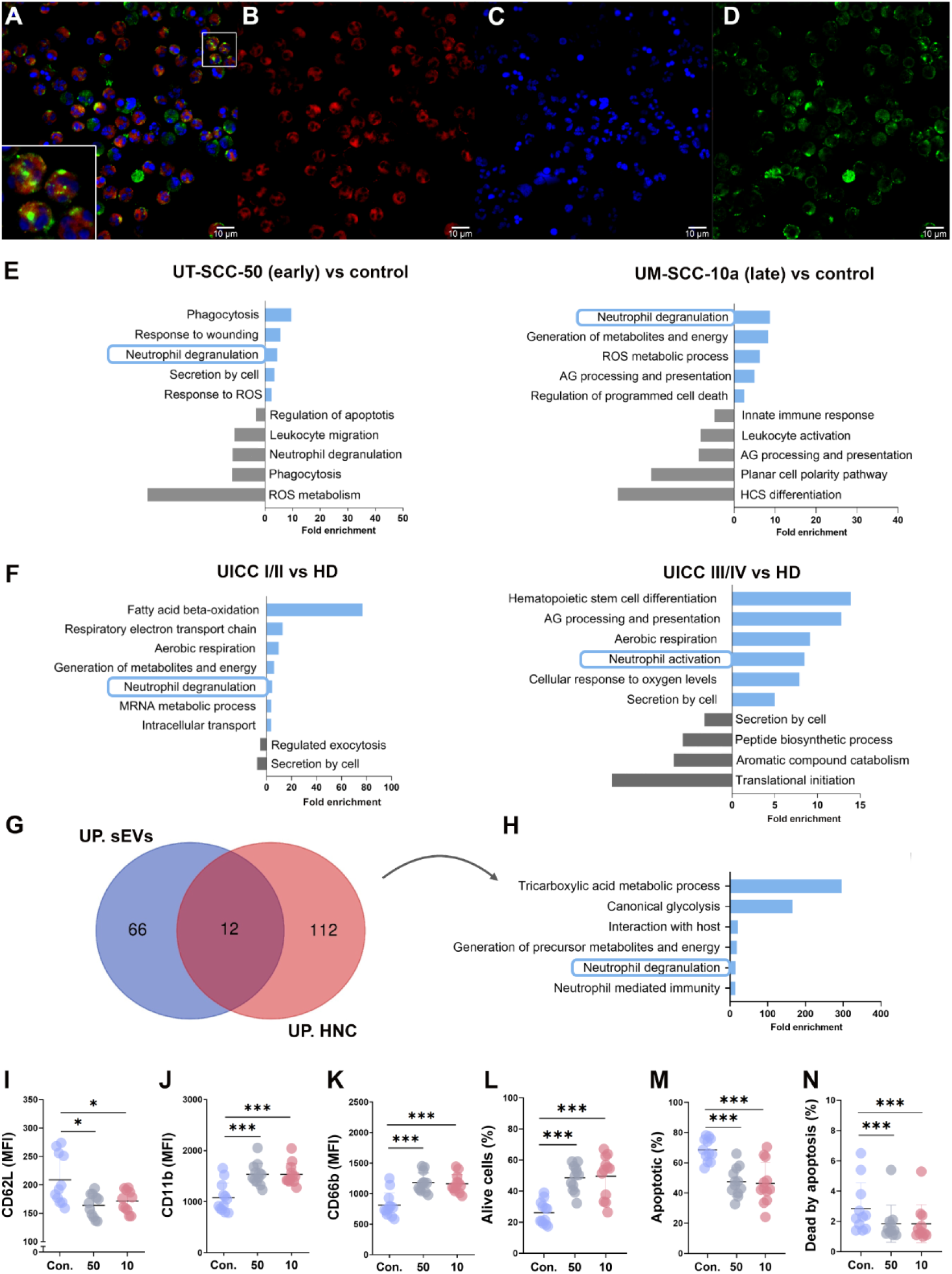
Tumor-derived sEVs are efficiently internalized by neutrophils and repolarized them. The ability of neutrophils to internalize tumor-derived sEVs was investigated by examining the uptake of sEVs isolated using size exclusion chromatography (SEC) (**A-D**). Neutrophils were co-incubated together with DID to label the cell surface (**B**) and PureBlu Hoechst 33342 to visualize the DNA (**C**). CFSE dye was used to visualize sEVs (**D**); (**A**) merged. To eliminate unbound CFSE dye, we employed SEC as described previously. Scale bar 10 µm. Comparative analysis of proteome enrichment in neutrophils educated by td-sEVs and neutrophils from HNC cancer patients demonstrated regulation of neutrophil degranulation by sEVs (E-H). (**E**) Fold enrichment of biological processes in neutrophils treated by sEVs from early-stage (UT-SCC-50) and late-stage (UM-SCC-10a) head and neck cancer (HNC) cell lines compared to healthy donors’ neutrophils (control). (**F**) Fold enrichment of biological processes in neutrophils Isolated from early-stage (UICC I/II) and late-stage (UICC III/IV) head and neck cancer patients compared to healthy donors (HD). The Venn diagram (**G**) shows the overlap between proteins upregulated in neutrophils from HNC patients (red circle) and those upregulated in neutrophils exposed to tumor-derived sEVs (blue circle), with a significance threshold of p<0.1. The intersection of the circles reveals 12 proteins that are similarly upregulated in both conditions, suggesting a common pathway activation. The bar graph (**H**) indicates the fold enrichment of biological processes associated with the commonly upregulated 12 proteins. Tumor-derived sEVs promote neutrophil survival and pro-tumoral phenotype (**I-N**). Freshly isolated human neutrophils after in vitro treatment with td-sEVS elevated cell activation, characterized by the downregulation of CD62L (I) and an increased expression of CD11b (J) and CD66b K). Expression of CD62L, CD11b, and CD66b on neutrophils was estimated by flow cytometry (**I-K**). Representative results from three replicate experiments are shown for (I-K), n=11 for Con. (control, HD neutrophils), n=11 for 50 (HD neutrophils treated by UM-SCC-50 EVs), n=11 for 10 (HD neutrophils treated by UT-SCC-10a EVs). The percentages of alive (L) (annexin V–/7-AAD–), apoptotic (**M**) (annexin V+ /7-AAD-), and dead by apoptosis (**N**) (annexin V+/7-AAD+) neutrophils after education with td-EVs were estimated by flow cytometry, n=12 for Con. (HD neutrophils), n=12 for 50 (HD neutrophils treated by UM-SCC-50 EVs), n=12 for 10 (HD neutrophils treated by UT-SCC-10a EVs). For comparison of two dependent groups Wilcoxon test was used. Data are shown as individual values with a median. * p < 0.05, *** p < 0.005

Furthermore, neutrophils treated by td-sEVs derived from early-stage (UT-SCC-50) and late-stage (UM-SCC-10a) HNC cell lines exhibited comparable pathway enrichment profiles (**Figure 4 E**) to those found in neutrophils isolated directly from early- and late-stage HNC patients (**Figure 4 F, Table S4**), with the prominent upregulation of proteins involved in neutrophil activation and degranulation processes.

In a detailed comparative proteomic analysis, we identified a range of proteins that were commonly upregulated both; in neutrophils isolated from the peripheral blood of HNC patients (denoted by the red circle) and those upregulated in neutrophils educated by td-sEVs (denoted by the blue circle) (**Figure 4 G**). Notably, many of these upregulated proteins are involved in crucial processes, such as neutrophil activation and degranulation (**Figure 4 H**). These findings indicate that the activation of neutrophils *in vitro* by td-sEVs mirrors the proteomic alterations observed in neutrophils from HNC patients.

In line with the proteomics data, we observed alterations of the neutrophil phenotype following *in vitro* education with td-sEVs, indicating elevated cell activation and degranulation, namely downregulation of CD62L (**Figure 4 I**) an increased expression of CD11b (**Figure 4 J**), and CD66b (**Figure 4 K**). Next, we assessed the percentages of alive (**Figure 4 L**) (annexin V^−^/7-AAD^−^), apoptotic (**Figure 4 M**) (early apoptosis, annexin V^+^ /7-AAD^−^) and dead by apoptosis (**Figure 4 N**) (late apoptosis, annexin V^+^/7-AAD^+^) neutrophils after education with td-sEVs. In agreement with the observed prolonged survival of pro-tumoral neutrophils in cancer patients, education with td-sEVs significantly prolonged survival of neutrophils and reduced their apoptosis (**Figure 4L-M, Sup.** Figure 5).

To exclude the possibility that soluble tumor-derived factors, rather than td-sEVs, are responsible for the observed phenomenon, healthy neutrophils were treated in parallel by sEVs depleted tumor-derived supernatants. Importantly, no notable alterations in marker expression were detected (**Sup. Figure S6**). The obtained results demonstrate that the changes observed in neutrophil phenotype and activity during cancer progression can be attributed to the internalization of td-sEVs, rather than to soluble factors in plasma and tumor microenvironment.

### Serpins associated with td-sEVs promote epithelial-to-mesenchymal transition (EMT) through neutrophil reprogramming

We observed that plasma-derived sEVs from HNC patients, in contrast to sEVs from healthy volunteers, potentiate tumor-supportive properties of neutrophils. Such educated neutrophils significantly promote survival of the tumor cells *in vitro* (**Figure 5 A-E**). Next we aimed to evaluate if neutrophils educated by tumor-derived sEVs change their functional capacities.

**Figure 5.**
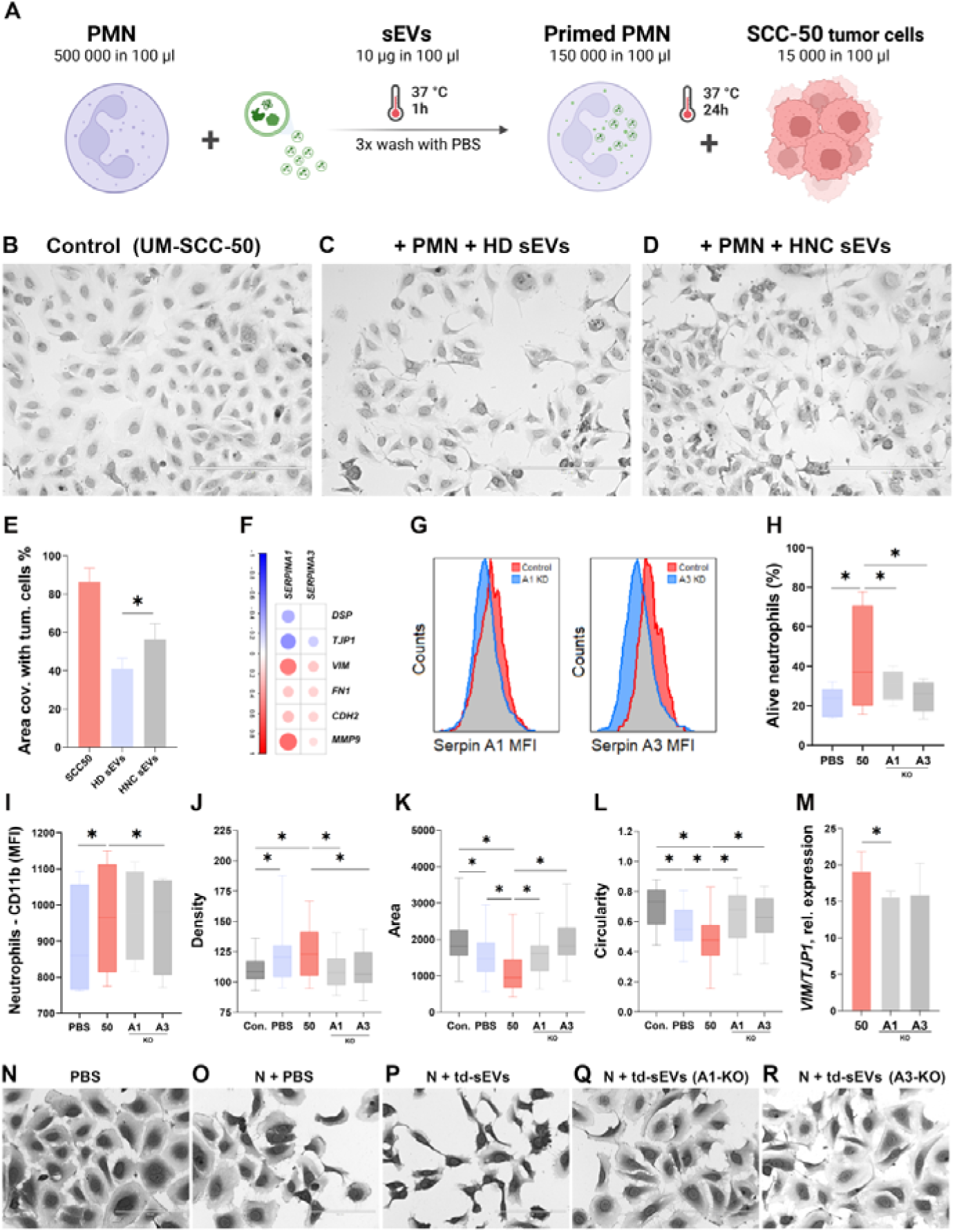
Tumor-derived sEVs with serpin cargo induce EMT through neutrophil reprogramming. (**A**) Scheme of experiment. (**B-D**) Representative images show tumor cell survival following co-culture with control neutrophils (**B**), neutrophils primed with healthy donor sEVs (**C**), and neutrophils primed with HNC- derived sEVs (**D**), scale bar 400µm. Neutrophils educated by HNC sEVs significantly enhanced tumor cell survival (E). Plot (F) shows a correlation analysis between serpin expression (A1, A3) in HNC tissues against EMT gene signatures. Serpin expression was negatively correlated with epithelial genes (DSP, TJP1) and positively correlated with mesenchymal markers (VIM, FN1, CDH2, MMP9), indicating EMT involvement. (**G**) CRISPR/Cas knockout (KO) of SERPIN A1 and A3 in UM-SCC-10a cells was verified by flow cytometry. (**H**) Reduced neutrophil viability after priming with serpin-deficient td-sEVs. (**I**) Downregulation of CD11b expression in neutrophils after priming with serpin-deficient td-sEVs. Plots (J-L) show results of morphological analysis of tumor cells co-cultured with neutrophils primed with td-sEVs from serpin-sufficient or serpin-KO cells; the analysis revealed significant differences in cell density (**J**), area (**K**), and circularity (**L**), indicating mesenchymal transformation. Neutrophils educated with serpin-deficient td-sEVs failed to induce EMT, as indicated by decreased ratio of VIM to TJP1 (**M**) in tumor cells. (**N-R**) Representative images of tumor cells co-incubated with neutrophils primed with td-sEVs from serpin-sufficient and -KO cells show distinct changes in cellular morphology, consistent with EMT, scale bar 200µm. For comparison of two dependent groups Wilcoxon test was used. Data are shown as bars with a median. Correlations were analyzed with the Spearman R test. * p < 0.05

As expression of serpins (A3 and A1) in tumor tissue correlated with the gene signature of EMT (negative correlation with epithelial genes *DSP* and *TJP*, and positive correlation with a set mesenchymal genes *VIM*, *FN1*, *CDH2* and *MMP9*) in HNC (**Figure 5 F**), we assumed that serpin cargo of td-sEVs stimulated EMT-activatory capacity of neutrophils. To test if serpin cargo of tumor-derived sEVs is responsible for the observed reprogramming of neutrophils, we performed CRISPR/Cas serpin-KO in UM-SCC-10a cell line and compared the effects of sEVs from such cell lines with serpin-sufficient sEVs serpin-KO. Downregulated expression of serpins in KO tumor cells versus WT cells was verified with flow cytometry (**Figure 5 G**). Next, we educated neutrophils with td-sEVs isolated from both: serpin-deficient and -sufficient cell lines, and assessed their phenotype and functions. Importantly, depletion of both serpins in sEVs resulted in decreased neutrophil viability (**Figure 5 H**). We also observed decreased CD11b expression in neutrophils primed with serpin-depleted td-sEVs (**Figure 5 I**).

Co-incubation of td-sEVs-primed neutrophils vs control neutrophils with tumor cells (scheme of experiment **Figure 5 A**) resulted in the apparent change of tumor cell phenotype towards mesenchymal, demonstrated by the analysis of cellular morphology (density, area, circularity) (**Figure 5 J-L, N-R**). Importantly, neutrophils educated with serpin-deficient td-sEVs failed to induce EMT, as judged by the relative expression of VIM to TJP (**Figure 5 M**).

### Serpins associated with td-sEVs promote immunosuppression through neutrophil reprogramming

As neutrophils play an important immunoregulatory role during tumor progression, we aimed to evaluate how tumor-derived sEVs modulate their immunoregulatory properties (**scheme of experiment Figure 6A**). Indeed, we could demonstrate the significant reduction of CD8+ T cell killing capacity after co-incubation with neutrophils primed previously by HNC patient-derived plasma sEVs, in comparison to sEVs from healthy donors (**Figure 6 B - F**).

**Figure 6.**
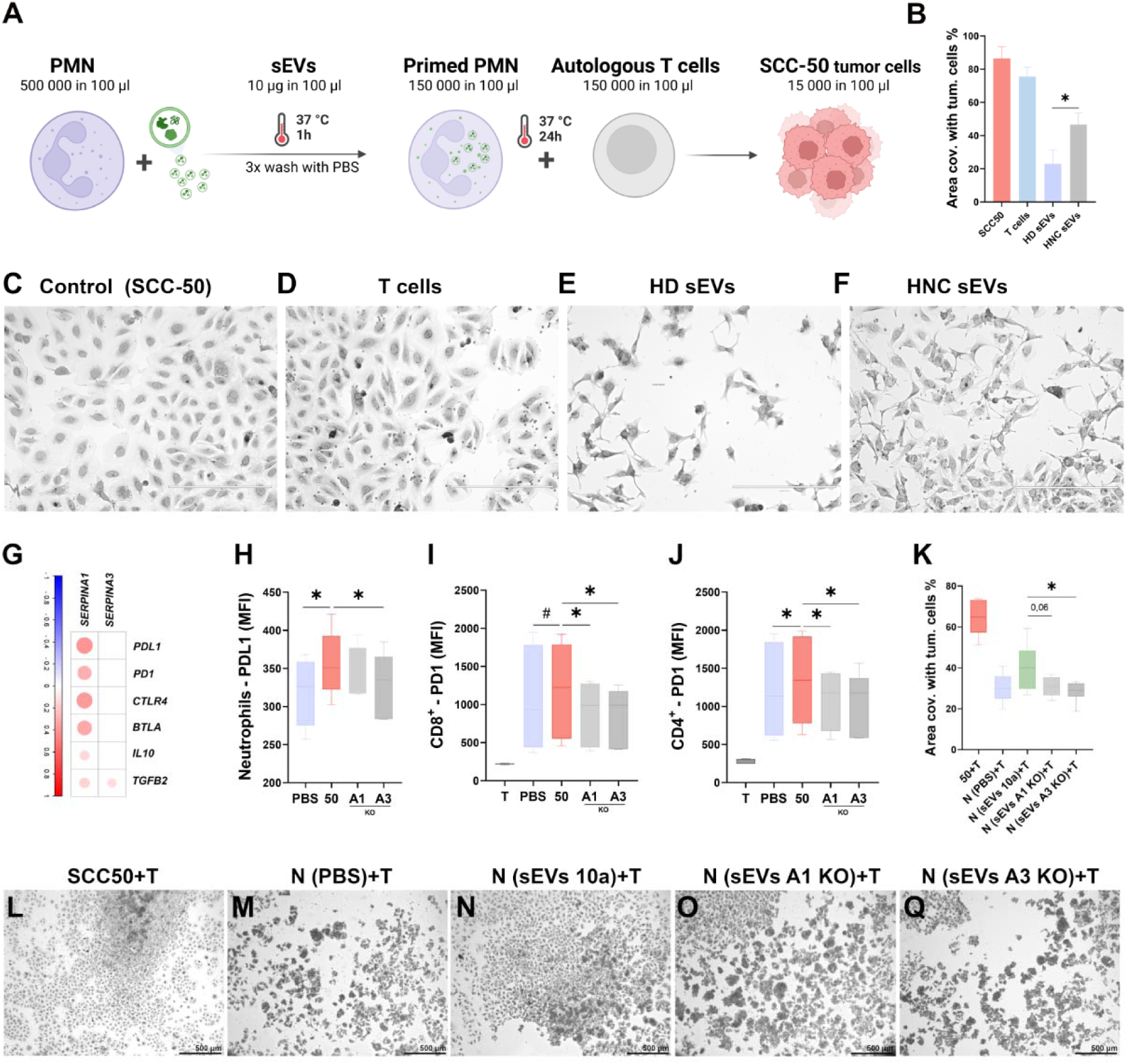
Tumor-derived sEVs with serpins cargo suppress T cells via neutrophil reprogramming. (**A**) Scheme of an experiment. (**B-F**) Tumor cell survival following co-incubation with T cells and neutrophils primed by healthy donor (HD) or HNC patient-derived sEVs, quantification (B) and representative images (C-F), scale bar 400µm. Neutrophils primed with HNC sEVs significantly reduced T cell killing potential compared to those primed with HD sEVs. (**G**) Expression of serpins (A1, A3) in HNC tumor tissue correlates with immunosuppressive gene signatures (PDL1, PD1, CTLA4, BTLA, IL10, TGFß2). (**H**) Flow cytometry showed increased PDL1 expression on neutrophils primed with td-sEVs from serpin-sufficient cells in comparison to serpin-deficient sEVs. (**I-J**) Education of neutrophils with serpin-deficient sEVs improves the capacity of neutrophils to stimulate T cells, as measured by decreased expression of PD-1 on CD8^+^ and CD4^+^ T cells. (**K-P**) Neutrophils primed by serpin-deficient sEVs restored T cell killing potential against tumor cells, while those primed by serpin-sufficient sEVs led to increased tumor survival, quantification (**K**) and representative images (**L-Q**), scale bar 500 µm. For comparison of two dependent groups Wilcoxon test was used. Data are shown as bars with a median. Correlations were analyzed with the Spearman R test. * p < 0.05.

Analysis of the open assess GEO database GSE65858^28^ demonstrated the significant correlation between the expression of serpins (A1 and A3) in tumor tissue with the gene signature of immunosuppression (PDL1, PD1, CTLA4, BTLA, IL10, TGFβ2) in HNC (**Figure 6 G**). To validate the contribution of td-sEVs-associated serpins in neutrophil-mediated immunosuppression, we evaluated the capacity of td-sEVs-primed neutrophils to induce immunosuppressive state of T cells, but also tumor cells in the co-culture experiments. First, we demonstrated the induction of PDL1 expression on neutrophils primed by td-sEVs, which was significantly higher in case of WT sEVs, compared to serpin-deficient sEVs (**Figure 6 H**). Importantly, co-incubation of such treated neutrophils with T cells in the presence of tumor cells induced expression of PD1 on both: CD8+ and CD4+ T cells (**Figure 6 I, J**). Moreover, serpin-deficient sEVs rescue neutrophils’ immunostimulatory properties, as autologous T cells primed by such neutrophils demonstrated elevated killing potential against tumor cells (**Figure 6 K**), in comparison T cells primed by neutrophils after co-incubation with WT sEVs (**Figure 6 L-Q**).

## Discussion

Extracellular vesicles, together with cytokines or growth factors have been shown to influence the activity of neutrophils in cancer. Here we uncover a previously unknown mechanism by which tumor-derived sEVs, through their serpin cargo, can reprogram neutrophils to adopt a tumor-supporting and immunosuppressive phenotype.

Tumors exploit sEVs as critical mediators of immunosuppression by directly inhibiting effector immune cells and promoting immunosuppressive cell populations, modulating immune checkpoints, and impeding lymphocyte infiltration into the tumor site^42^. In td-sEVs cargo, the following molecules were previously detected to directly influence the functions of immune cells: PDL1, PD1, CTLA4, FasL, TGFß and IL-10^43^.

We identified serpins as a component of tumor-derived sEVs, which is responsible for neutrophil reprogramming. The **SER**ine **P**rotease **IN**hibitor (serpin) superfamily is a diverse group of irreversible protease inhibitors. Despite the significant diversity of serpins, they can be grouped according to their structure or functions, e.g., inhibitors of coagulation, inhibitors of proteinases, etc. Nevertheless, the role of human serpins in a variety of biological processes is not fully understood^44^. Hereinafter, when referring to the term serpin, we mean SERPIN A1 and SERPIN A3, unless otherwise indicated.

Both; SERPIN A1, also known as alpha-1 antitrypsin, and SERPIN A3 - alpha-1 antichymotrypsin, belong to the group of inhibitors of proteinases with the main function of maintaining cellular homeostasis. Their function is crucial for controlling protease activity during inflammation to prevent tissue damage^45,46^. In the context of cancer, protease inhibition plays an important role in supporting tumor cell survival and spread. Elevation of SERPIN A1 in plasma has been observed in the course of a large number of malignant diseases like hepatocellular carcinoma^47^, colorectal cancer^48^, prostate and lung cancer^49^, bladder urothelial carcinoma and stomach adenocarcinoma^50^. Moreover, upregulation of several types of serpins by the tumor tissue had been previously demonstrated to be associated with insufficient response to therapies and poor prognosis^51^. These findings align closely with our observations regarding serpin cargo within plasma-derived extracellular vesicles (pd-sEVs) obtained from HNC patients. We found a significant elevation in serpin content already at the initial cancer stages, followed by a constant increase during tumor progression. This shows a correlation between the expression levels of serpins in pd-sEVs and the severity of the disease, making serpin cargo of plasma sEVs a promising biomarker in HNC.

Tumors upregulate the expression of serpins for several reasons, primarily related to enhancing their survival, immune evasion, and migration, thus promoting metastasis^52–54^. Serpins also reprogram immune cells to become immunosuppressive. Inhibition of serine proteases by serpins was demonstrated to induce an alternatively activated (M2) macrophage phenotype, which is associated with anti-inflammatory and tissue repair functions^55^. High levels of SERPIN A3 are associated with immune suppression in glioma, affecting macrophage infiltration and function^56^. On the contrary, loss of certain serpins leads to a decrease in tumor-promoting M2-like macrophages and associated cytokines like CCL2, which are crucial for tumor metastasis^57^. In the autosomal co-dominant genetic defect of SERPIN A1 called alpha-1 antitrypsin deficiency, unresolved inflammation and tissue damage due to macrophage dysfunction can be observed^58^. Serpins such as SERPIN B9 can inhibit granzyme B (GrB), a protease involved in inducing apoptosis in target cells^59^, which in the context of cancer can lead to the resistance against CTL-mediated killing, thereby promoting immune evasion and tumor survival^60^. In the context of neutrophils, Serpins A1 and A3 are known to inhibit neutrophil elastase (NE) and cathepsin G, respectively^41^.

Serpins play a pivotal role in the inhibition of apoptosis, thereby promoting cell survival. Studies have highlighted specific serpins, including Serpin A3, Serpin B1, and Serpin B9, as key players in enhancing cell survival^61^. These serpins exert their effects by inhibiting proteases involved in apoptotic signaling pathways, such as granzymes or caspases^62,63^. This helps to maintain a reserve of neutrophils in the bone marrow^64^. These findings are consistent with the prolonged lifespan of pro-tumoral neutrophils that we observe in cancer patients. In agreement with this, we observed that the exposure to td-sEVs significantly prolonged the survival of neutrophils and reduced their apoptosis. These results provide further support for the role of serpins in promoting cell survival, particularly in the context of cancer progression.

We further demonstrated the activated phenotype (low expression of CD62L) and elevated expression of CD11b and CD66b on neutrophils educated with serpin-containing td-sEVs, which is a sign of elevated degranulation^65^. Such increased degranulation could be a compensatory mechanism to overcome the inhibition of released enzymes. The observed phenomenon requires further investigation.

Besides limiting the toxicity of neutrophil-derived enzymes, serpins inhibit the cytotoxicity of neutrophils against tumor cells. Thus, NE was reported to kill cancer cells selectively and attenuate tumor development^66^. Similarly, Neutrophil cathepsin G facilitates neutrophil anti-tumor cytotoxicity^67^. Data presented by Kudo^68^ suggests that neutrophil-derived cathepsin G prevents tumor cell invasion by inducing tight cell-cell adhesion. Although we did not observe changes in the killing of tumor cells by td-sEVs-primed neutrophils, we could show elevated epithelial-mesenchymal transition upon stimulation by td-sEVs-primed neutrophils, which was dependent on the presence of serpins.

As we previously demonstrated, neutrophils play a significant role in the regulation of adaptive immune responses in cancer by presenting antigens and activating lymphocytes, or by expressing immune checkpoint molecules such as PD-L1 and induction of T cell dysfunctional state^6,69,70^. As previously shown by us and others, T-cell exhaustion is a hallmark of oncoimmunology and a main avenue of cancer-induced immunosuppression^71^. T cell states seem to go through different states of dysfunction, which lie on a continuum^72^. The stage of terminal dysfunction is characterized by the high expression PD1 as well as multiple other checkpoint receptors, such as CD39, Tim-3, TIGIT^73^. Importantly, reviving dysfunctional T cells is the main treatment modality of currently approved immunotherapy agents in HNSCC, namely anti-PD1 antibodies^71^.

In this current study, we show a new mechanism of inducing T cell dysfunction: the td-sEVs-associated serpins induced neutrophil-mediated upregulation of checkpoint receptors in effector T cells. This also impaired the function of these T cells, as demonstrated by the reduced T cell-mediated tumor cell killing. This might have therapeutic implications for the current immunotherapy treatment strategy and could inform future combination therapies. This finding might have therapeutic implications for the future immunotherapy treatment strategies.

Although the exact mechanism of how EV-associated serpins contribute to neutrophil-mediated T cell dysfunction is unknown, the interaction of serpins with neutrophil granule enzymes might be responsible for this phenomenon. While NE was demonstrated to stimulate adaptive immunity via the increased peptide presentation and downregulation of T cell inhibitory molecules^74^, cathepsin G was shown to be important for antigen processing and presentation^75^. Serpins A1 and A3 by inhibiting NE and cathepsin G, respectively, might contribute to the tumor-induced suppression of adaptive immunity. In agreement, our data showed that selective knock-down of serpins in sEVs reprogrammed neutrophils to potentiate anti-tumor T cell responses.

Therapeutic repolarization of neutrophils into anti-tumor effectors remains an attractive perspective in the cancer treatment. Since serpin-enriched td-sEVs stimulate pro-tumor polarization of neutrophils, various therapeutic approaches might be potentially applied to minimize this negative impact, including targeting serpins or td-sEVs release or uptake. As overexpression of serpins was proven to be a prognostic and predictive marker in a variety of malignancies, they represent an important drug target for treating cancer. Nevertheless, at the current point, the direct inhibition of serpins remains challenging. Several miRNA are known to directly interfere with serpins, and few chemical compounds were described to inhibit serpins activity or decrease their expression^76^. Targeting small extracellular vesicles as a therapeutic strategy in cancer has gained significant attention due to their role in intercellular communication and their potential as drug delivery vehicles^77^. Different approaches, such as prevention of sEV biogenesis and release by tumor cells (e.g. with inhibition of endosomal sorting complex required for transport or neutral sphingomyelinase), removal of EVs from the entire circulatory system with an affinity plasmapheresis platform, as well as inhibition of EV internalization by recipient cells mediated by phosphatidylserine, heparan sulfate proteoglycans, tetraspanins and CD54 were also investigated and demonstrated promising results^78^. Based on our results, we propose that targeting serpins in EVs, in combination with immunotherapy, could provide an effective strategy to further improve antitumor immune responses.

## Conclusion

These findings underscore the critical role of td-sEVs-associated SERPIN A1 and A3 in reprogramming neutrophils, fostering a pro-tumoral environment, facilitating EMT of tumor cells, immune evasion, and promoting tumor progression. We further propose a new mechanism for neutrophil-mediated induction of dysfunctional T cells. Understanding these mechanisms opens new avenues for therapeutic strategies targeting sEVs-mediated neutrophil education in cancer.

## Acknowledgments

We gratefully acknowledge the Imaging Center Essen (IMCES) for equipment support. We acknowledge support by the Open Access Publication Fund of the University of Duisburg-Essen. This work was supported by the DFG (DFG/JA-2461/5-1 and 7-1), CRC TRR332 project A05 to JJ and DFG (Project-ID 418179183 – KFO 337 (HE 5294/2-1 (IH)). All illustrations were designed using BioRender. A.B. was supported by the scholarship NAWA PPN/STA/2021/1/00027

## Author contributions

*Conceptualization*: M.D., E.P., and J.J.; *methodology:* M.D., E.P., M.P., I.T., M.G, M.S., H.H., and J.J.; *software*: M.D., E.P., M.P.; *validation*: M.D., E.P., E.S., and J.J.; *formal analysis*: M.D., E.P., E.S.; *investigation*: M.D., E.P., E.S., N.K., A.B., D.S., A.Z., M.G., M.S., M.R., D.F., M.P.; *resources*: S. Lang, C.K., M.P., B.G., I.H., and J.J.; *data curation*: M.D, E.P., and J.J.; *writing – original draft preparation*: M.D., E.P.; *writing – review & editing*: E.S., A.B., D.S., M.G., M.S., M.R., D.F., I.O., I.H., B.T., M.P., C.K. S.L., and J.J.; *visualization*: M.D., E.P; *supervision*: J.J.; *project administration*: J.J.; *funding acquisition*: J.J.

---

**Figure S1.**
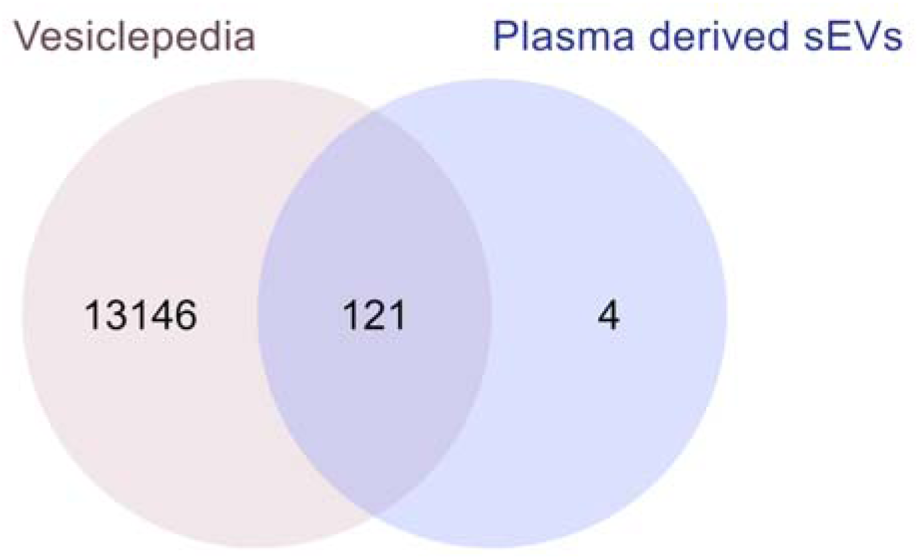
Protein content of sEVs. The analysis of the Vesiclepedia database revealed that 121 proteins from our list have been previously found to be associated with extracellular vesicles, with only 4 of them being identified in our study for the first time as a part of extracellular vesicles, and not previously annotated in this database. Data visualization was performed using FunRich (version 3.1.4) (http://www.funrich.org/).

**Figure S2.**
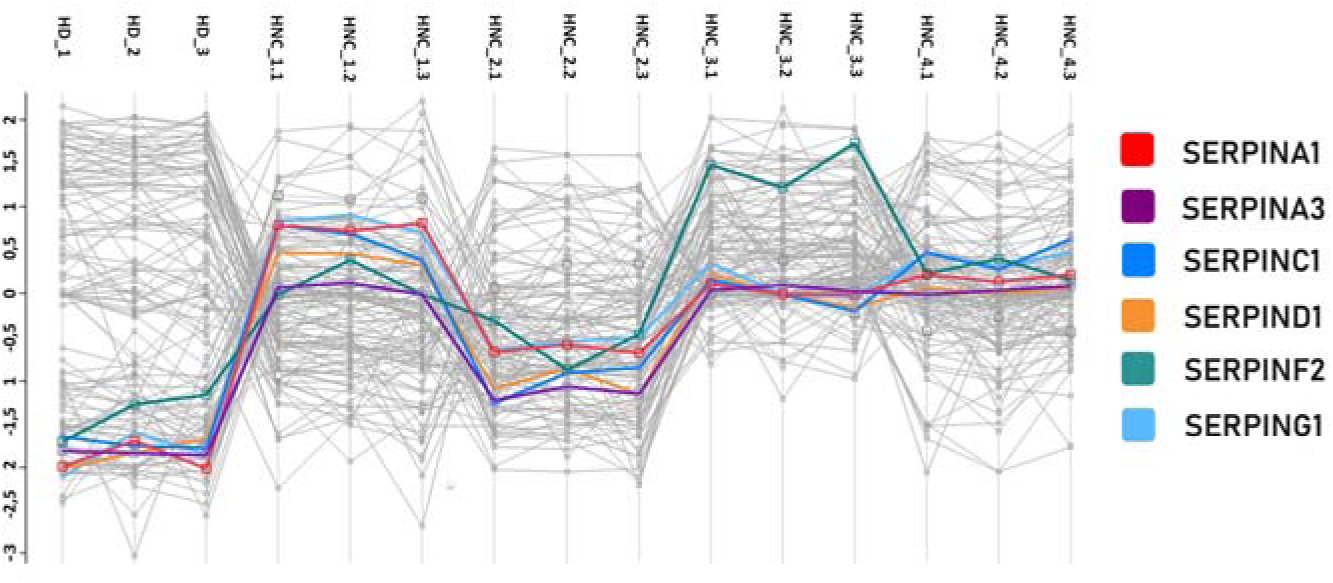
Serpin expression increases during cancer progression. Visualization of Z-score normalized Label-Free Quantification (LFQ) intensities for serpins (SERPINA1, SERPINA3, SERPINC1, SERPIND1, SERPINF2, SERPING) among healthy donors (HD) and patients with Head and Neck Cancer (HNC) at various stages (UICC I-IV). The y-axis on the left represents Z-score normalized LFQ intensity, with sample names labeled on the top. Each color-coded line corresponds to one of the mentioned serpins.

**Figure S3.**
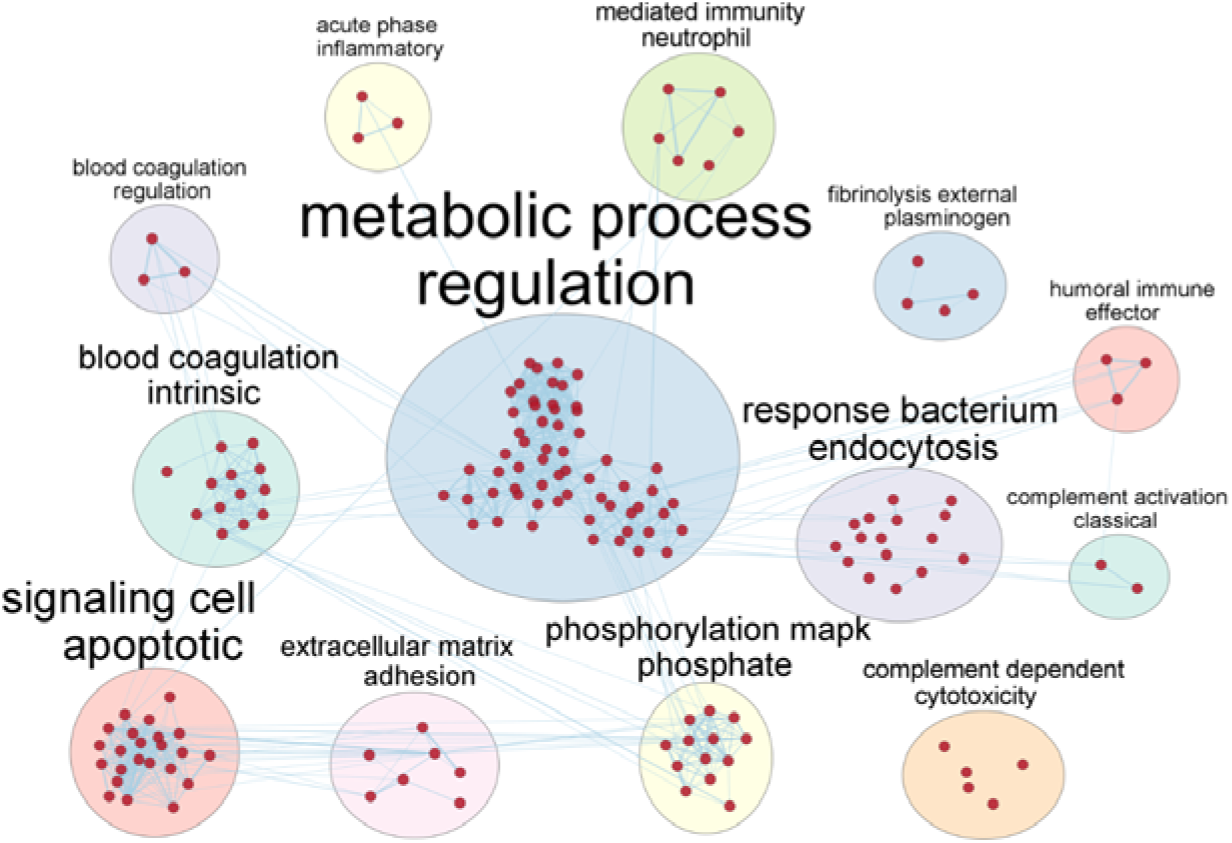
The proteome of sEVs isolated from the plasma of HNC patients identifies the regulation of multiple neutrophil-specific signaling pathways. Plasma-derived sEVs were isolated from individuals with HNC and proceeded for proteomics. Pathways enrichment analysis of identified proteins depicted as an enrichment map (q-value<0.0001, enrichment analysis performed in Enrichment Map app (https://apps.cytoscape.org/apps/enrichmentmap) by using Cytoscape (https://cytoscape.org/). The resulting enrichment map was annotated using AutoAnnotate (https://apps.cytoscape.org/apps/autoannotate) with standard settings with the aim of seeing the general overview of the functions of isolated proteins. In this format, large clusters are no more important than single nodes but it is easy to identify the major themes of analysis.

**Figure S4.**
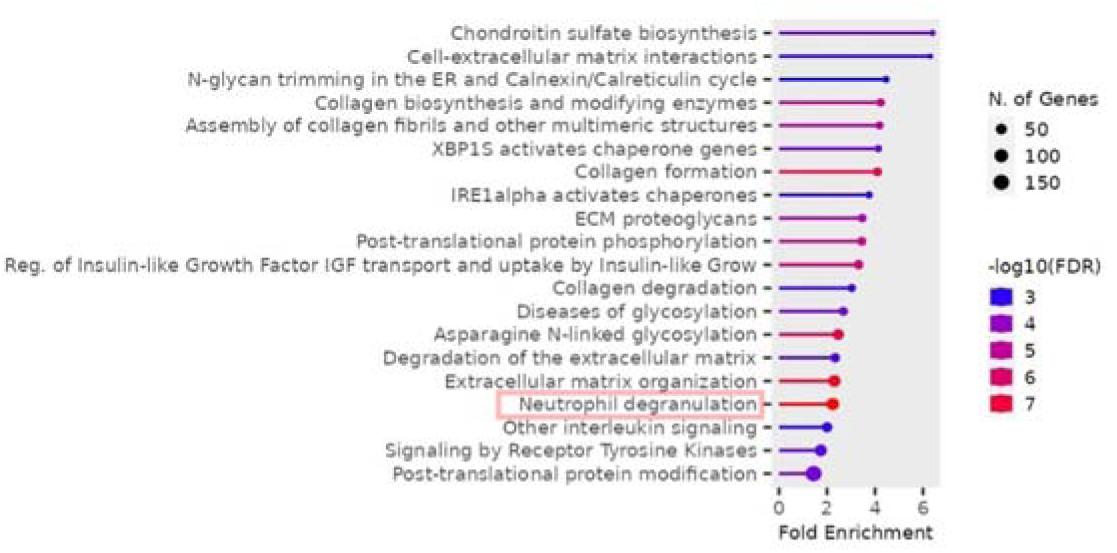
Neutrophil degranulation plays an important role in HNC progression. The publicly available database GSE65858 (https://www.ncbi.nlm.nih.gov/geo/query/acc.cgi?acc=GSE65858) was used to identify overrepresented pathways. Graphical visualization was performed using ShinyGO (http://bioinformatics.sdstate.edu/go/).

**Figure S5.**
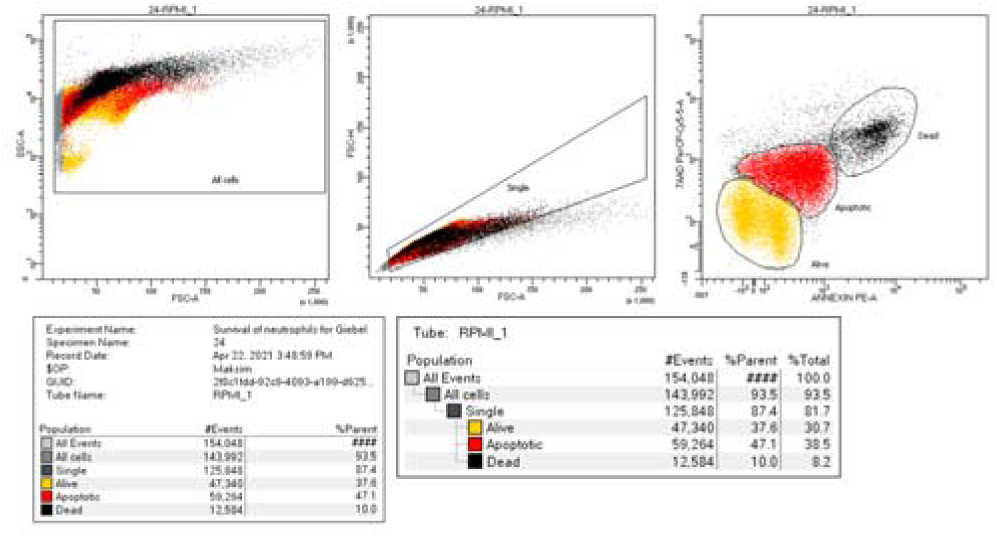
Gating strategy example. Survival assay. Neutrophils were gated into distinct populations based on their Annexin (PE) expression level and 7AAD (PerCP-Cy5). Cells characterized by high levels of 7AAD^high^ and Annexin^high^ were classified as dead cells, while those displaying moderate levels of both markers 7AAD^mid^ and Annexin^mid^ were designated as apoptotic. Cell exhibiting low levels of 7AAD^low^ and Annexin^low^ were categorized as alive.

**Figure S6.**
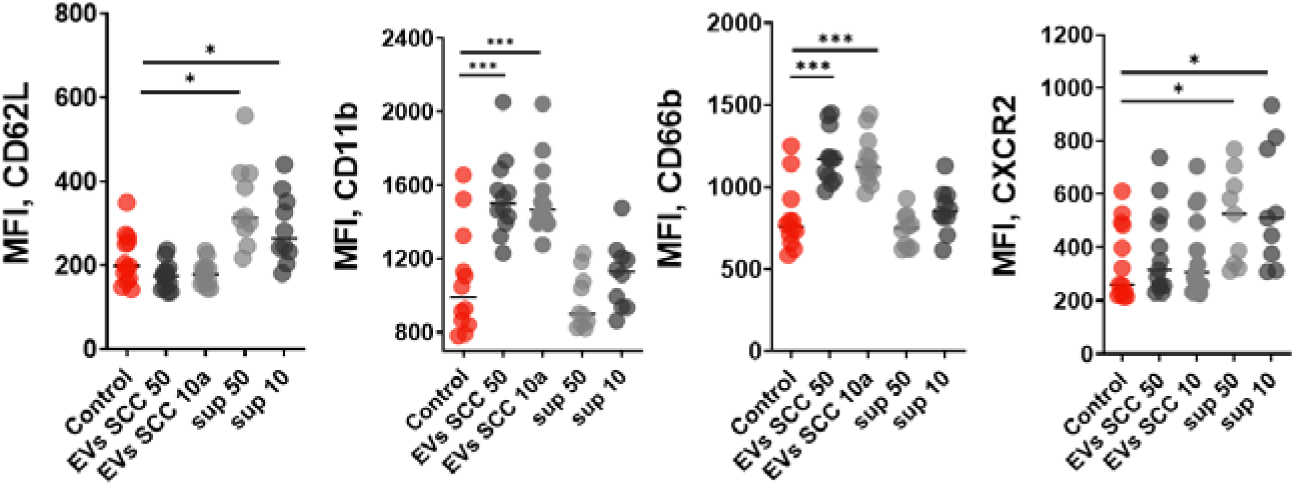
Changes in neutrophil behavior during cancer progression are linked to uptake of td-sEVs, not soluble factors. In order to rule out the possibility that soluble tumor-derived factors, rather than td-sEVs, are responsible for the observed phenomenon, healthy neutrophils were treated concurrently with both t - sEVs and td-sEVs -depleted tumor-derived supernatants. No significant alterations in marker expression were detected following incubation with td-sEVs -depleted tumor-derived supernatants. Marker’s expression (CD62L, CD11b, CD66b, CXCR2) on neutrophils was estimated by flow cytometry. Representative results from three replicate experiment are shown, n=13 for Control (HD neutrophils), n=13 for td-sEVs SCC50 (HD neutrophils treated by UM-SCC-50 sEVs), n=13 for td-sEVs SCC 10a (HD neutrophils treated by UT-SCC-10a td-sEVs), n=10 for sup 50 (EVs depleted UM-SCC-50 cell culture supernatant), n=10 for sup 10 (EVs depleted UM-SCC-10a cell culture supernatant). For comparison of two independent groups, Mann-Whitney U test was used, and for comparison of multiple independent groups Kruskal-Wallis (ANOVA) was used. Data are shown as median with interquartile range and individual values with median. Data are shown as an individual values with median. * p < 0.05, *** p < 0.005.

## References

1. Jablonska, J., Leschner, S., Westphal, K., Lienenklaus, S. & Weiss, S. Neutrophils responsive to endogenous IFN-beta regulate tumor angiogenesis and growth in a mouse tumor model. J Clin.Invest 120, 1151–1164 (2010).

2. Fridlender, Z. G. et al. Polarization of tumor-associated neutrophil phenotype by TGF-beta: ‘N1’ versus ‘N2’ TAN 1. Cancer Cell 16, 183–194 (2009).

3. Andzinski, L. et al. Type I IFNs induce anti-tumor polarization of tumor associated neutrophils in mice and human. Int.J.Cancer 138, 1982–1993 (2016).

4. Decker, A. S. et al. Prognostic Role of Blood NETosis in the Progression of Head and Neck Cancer. Cells 8, (2019).

5. Bordbari, S. et al. SIRT1-mediated deacetylation of FOXO3a transcription factor supports pro-angiogenic activity of interferon-deficient tumor-associated neutrophils. Int J Cancer 150, 1198–1211 (2022).

6. Pylaeva, E. et al. During early stages of cancer, neutrophils initiate anti-tumor immune responses in tumor-draining lymph nodes. Cell Rep 40, (2022).

7. Trellakis, S. et al. Peripheral blood neutrophil granulocytes from patients with head and neck squamous cell carcinoma functionally differ from their counterparts in healthy donors. Int J Immunopathol Pharmacol 24, 683–693 (2011).

8. Albrengues, J. et al. Neutrophil extracellular traps produced during inflammation awaken dormant cancer cells in mice. Science 361, (2018).

9. Anselmi, M. et al. Melanoma Stem Cells Educate Neutrophils to Support Cancer Progression. Cancers (Basel) 14, (2022).

10. Wu, C. F. et al. The lack of type I interferon induces neutrophil-mediated pre-metastatic niche formation in the mouse lung. Int.J.Cancer (2015).

11. Van Niel, G., D’Angelo, G. & Raposo, G. Shedding light on the cell biology of extracellular vesicles. Nat Rev Mol Cell Biol 19, 213–228 (2018).

12. Tengler, L. et al. Plasma-derived small extracellular vesicles unleash the angiogenic potential in head and neck cancer patients. Mol Med 29, (2023).

13. Zitvogel, L. et al. Eradication of established murine tumors using a novel cell-free vaccine: dendritic cell-derived exosomes. Nat Med 4, 594–600 (1998).

14. Skokos, D. et al. Mast cell-derived exosomes induce phenotypic and functional maturation of dendritic cells and elicit specific immune responses in vivo. J Immunol 170, 3037–3045 (2003).

15. Raposo, G. et al. B lymphocytes secrete antigen-presenting vesicles. J Exp Med 183, 1161–1172 (1996).

16. Jablonska, J. et al. Evaluation of Immunoregulatory Biomarkers on Plasma Small Extracellular Vesicles for Disease Progression and Early Therapeutic Response in Head and Neck Cancer. Cells 11, (2022).

17. Gehrmann, U. et al. Synergistic induction of adaptive antitumor immunity by codelivery of antigen with α-galactosylceramide on exosomes. Cancer Res 73, 3865–3876 (2013).

18. Hoffmann, T. K. et al. Alterations in the p53 pathway and their association with radio- and chemosensitivity in head and neck squamous cell carcinoma. Oral Oncol 44, 1100–1109 (2008).

19. Lin, C. J. et al. Head and neck squamous cell carcinoma cell lines: established models and rationale for selection. Head Neck 29, 163–188 (2007).

20. Bradford, C. R. et al. Human papillomavirus DNA sequences in cell lines derived from head and neck squamous cell carcinomas. Otolaryngol Head Neck Surg 104, 303–310 (1991).

21. Baker, S. R. An in vivo model for squamous cell carcinoma of the head and neck. Laryngoscope 95, 43– 56 (1985).

22. Virts, E. L. et al. AluY-mediated germline deletion, duplication and somatic stem cell reversion in UBE2T defines a new subtype of Fanconi anemia. Hum Mol Genet 24, 5093–5108 (2015).

23. Wiek, C. et al. Identification of amino acid determinants in CYP4B1 for optimal catalytic processing of 4-ipomeanol. Biochem J 465, 103–114 (2015).

24. Wisniewski, J. R. & Gaugaz, F. Z. Fast and sensitive total protein and peptide assays for proteomic analysis. Anal Chem 87, 4110–4116 (2015).

25. Rappsilber, J., Ishihama, Y. & Mann, M. Stop And Go Extraction tips for matrix-assisted laser desorption/ionization, nanoelectrospray, and LC/MS sample pretreatment in proteomics. Anal Chem 75, 663–670 (2003).

26. Perez-Riverol, Y. et al. The PRIDE database resources in 2022: a hub for mass spectrometry-based proteomics evidences. Nucleic Acids Res 50, D543–D552 (2022).

27. Théry, C., Amigorena, S., Raposo, G. & Clayton, A. Isolation and Characterization of Exosomes from Cell Culture Supernatants and Biological Fluids. Curr Protoc Cell Biol 30, 1–29 (2006).

28. Wichmann, G. et al. The role of HPV RNA transcription, immune response-related gene expression and disruptive TP53 mutations in diagnostic and prognostic profiling of head and neck cancer. Int J Cancer 137, 2846–2857 (2015).

29. Shannon, P. et al. Cytoscape: a software environment for integrated models of biomolecular interaction networks. Genome Res 13, 2498–2504 (2003).

30. Merico, D., Isserlin, R., Stueker, O., Emili, A. & Bader, G. D. Enrichment map: a network-based method for gene-set enrichment visualization and interpretation. PLoS One 5, (2010).

31. Ge, S. X., Jung, D., Jung, D. & Yao, R. ShinyGO: a graphical gene-set enrichment tool for animals and plants. Bioinformatics 36, 2628–2629 (2020).

32. Goedhart, J. & Luijsterburg, M. S. VolcaNoseR is a web app for creating, exploring, labeling and sharing volcano plots. Sci Rep 10, (2020).

33. Tang, D. et al. SRplot: A free online platform for data visualization and graphing. PLoS One 18, (2023).

34. Pathan, M. et al. FunRich: An open access standalone functional enrichment and interaction network analysis tool. Proteomics 15, 2597–2601 (2015).

35. Pathan, M. et al. A novel community driven software for functional enrichment analysis of extracellular vesicles data. J Extracell Vesicles 6, (2017).

36. Hong, C. S., Funk, S. & Whiteside, T. L. Isolation of Biologically Active Exosomes from Plasma of Patients with Cancer. Methods in molecular biology (Clifton, N.J.) 1633, 257–265 (2017).

37. Ludwig, N. et al. Isolation and Analysis of Tumor-Derived Exosomes. Current Protocols in Immunology 127, (2019).

38. Hong, C. S., Funk, S., Muller, L., Boyiadzis, M. & Whiteside, T. L. Isolation of biologically active and morphologically intact exosomes from plasma of patients with cancer. J.Extracell.Vesicles. 5, 29289 (2016).

39. Welsh, J. A. et al. Minimal information for studies of extracellular vesicles (MISEV2023): From basic to advanced approaches. Journal of extracellular vesicles 13, (2024).

40. Lecot, P. et al. Gene signature of circulating platelet-bound neutrophils is associated with poor prognosis in cancer patients. International journal of cancer 151, 138–152 (2022).

41. Johnston, A. et al. IL-1 and IL-36 are dominant cytokines in generalized pustular psoriasis. The Journal of allergy and clinical immunology 140, 109–120 (2017).

42. Ahmadi, M., Abbasi, R. & Rezaie, J. Tumor immune escape: extracellular vesicles roles and therapeutics application. Cell communication and signalingfZ: CCS 22, (2024).

43. Ludwig, S. et al. Suppression of Lymphocyte Functions by Plasma Exosomes Correlates with Disease Activity in Patients with Head and Neck Cancer. Clin.Cancer Res. 23, 4843–4854 (2017).

44. Silverman, G. A. et al. The serpins are an expanding superfamily of structurally similar but functionally diverse proteins. Evolution, mechanism of inhibition, novel functions, and a revised nomenclature. The Journal of biological chemistry 276, 33293–33296 (2001).

45. Janciauskiene, S. et al. The Multifaceted Effects of Alpha1-Antitrypsin on Neutrophil Functions. Frontiers in pharmacology 9, 341 (2018).

46. de Mezer, M. et al. SERPINA3: Stimulator or Inhibitor of Pathological Changes. Biomedicines 11, (2023).

47. Comunale, M. A. et al. Linkage specific fucosylation of alpha-1-antitrypsin in liver cirrhosis and cancer patients: implications for a biomarker of hepatocellular carcinoma. PloS one 5, (2010).

48. Bujanda, L. et al. Evaluation of alpha 1-antitrypsin and the levels of mRNA expression of matrix metalloproteinase 7, urokinase type plasminogen activator receptor and COX-2 for the diagnosis of colorectal cancer. PloS one 8, (2013).

49. El-Akawi, Z. J., Al-Hindawi, F. K. & Bashir, N. A. Alpha-1 antitrypsin (alpha1-AT) plasma levels in lung, prostate and breast cancer patients. Neuro endocrinology letters 29, 482–484 (2008).

50. Kuai, X., Lv, J., Zhang, J., Xu, M. & Ji, J. Serpin Family A Member 1 Is Prognostic and Involved in Immunological Regulation in Human Cancers. International journal of molecular sciences 24, (2023).

51. Ibáñez-Molero, S. et al. SERPINB9 is commonly amplified and high expression in cancer cells correlates with poor immune checkpoint blockade response. Oncoimmunology 11, (2022).

52. Kwon, C. H. et al. Serpin peptidase inhibitor clade A member 1 is a biomarker of poor prognosis in gastric cancer. British Journal of Cancer 111, 1993 (2014).

53. Valiente, M. et al. Serpins promote cancer cell survival and vascular Co-option in brain metastasis. Cell 156, 1002–1016 (2014).

54. Kwon, C. H. et al. Snail and serpinA1 promote tumor progression and predict prognosis in colorectal cancer. Oncotarget 6, 20312 (2015).

55. Xu, N. et al. The Anti-Inflammatory Immune Response in Early Trichinella spiralis Intestinal Infection Depends on Serine Protease Inhibitor-Mediated Alternative Activation of Macrophages. Journal of immunology (Baltimore, Md.fZ: 1950) 206, 963–977 (2021).

56. Yuan, Q. et al. Highly expressed of SERPINA3 indicated poor prognosis and involved in immune suppression in glioma. Immunity, inflammation and disease 9, 1618–1630 (2021).

57. Smirnova, T. et al. Serpin E2 promotes breast cancer metastasis by remodeling the tumor matrix and polarizing tumor associated macrophages. Oncotarget 7, 82289–82304 (2016).

58. Lee, J. et al. Alpha 1 Antitrypsin-Deficient Macrophages Have Impaired Efferocytosis of Apoptotic Neutrophils. Frontiers in immunology 11, (2020).

59. Kaiserman, D. & Bird, P. I. Control of granzymes by serpins. Cell death and differentiation 17, 586–595 (2010).

60. Kimman, T. et al. Serpin B9 controls tumor cell killing by CAR T cells. Journal for immunotherapy of cancer 11, (2023).

61. Fehervari, Z. Superinduction! Nature immunology 18, 603 (2017).

62. Godovikova, V. & Ritchie, H. H. Dynamic processing of recombinant dentin sialoprotein-phosphophoryn protein. The Journal of biological chemistry 282, 31341–31348 (2007).

63. Liu, Y. N. et al. Renal retention of lipid microbubbles: a potential mechanism for flank discomfort during ultrasound contrast administration. Journal of the American Society of EchocardiographyfZ: official publication of the American Society of Echocardiography 26, 1474–1481 (2013).

64. Baumann, M., Pham, C. T. N. & Benarafa, C. SerpinB1 is critical for neutrophil survival through cell-autonomous inhibition of cathepsin G. Blood 121, 3900–3907 (2013).

65. Duerig, I. et al. Nonfunctional TGF-β/ALK1/ENG signaling pathway supports neutrophil proangiogenic activity in hereditary hemorrhagic telangiectasia. Journal of leukocyte biology 114, 639–650 (2023).

66. Cui, C. et al. Neutrophil elastase selectively kills cancer cells and attenuates tumorigenesis. Cell 184, 3163–3177.e21 (2021).

67. Sionov, R. V. et al. Neutrophil Cathepsin G and Tumor Cell RAGE Facilitate Neutrophil Anti-Tumor Cytotoxicity. Oncoimmunology 8, (2019).

68. Yui, S. et al. Cathepsin G, a neutrophil protease, induces compact cell-cell adhesion in MCF-7 human breast cancer cells. Mediators of inflammation 2009, (2009).

69. Hussain, T. et al. IFNAR1 Deficiency Impairs Immunostimulatory Properties of Neutrophils in Tumor-Draining Lymph Nodes. Frontiers in immunology 13, (2022).

70. Pylaeva, E. et al. B-Helper Neutrophils in Regional Lymph Nodes Correlate with Improved Prognosis in Patients with Head and Neck Cancer. Cancers 13, (2021).

71. Kürten, C. H. L. & Ferris, R. L. Neoadjuvant immunotherapy for head and neck squamous cell carcinoma. Laryngo-rhino-otologie 103, S167–S187 (2024).

72. Cillo, A. R. et al. Immune Landscape of Viral- and Carcinogen-Driven Head and Neck Cancer. Immunity 52, 183–199.e9 (2020).

73. Kürten, C. H. L. et al. Investigating immune and non-immune cell interactions in head and neck tumors by single-cell RNA sequencing. Nature communications 12, (2021).

74. Chawla, A., et al. Abstract 4843: Neutrophil elastase enhances adaptive immunity via increase of peptide presentation and downregulation of T cell inhibitory molecules. Cancer Research 74, 4843– 4843 (2014).

75. Burster, T., Macmillan, H., Hou, T., Boehm, B. O. & Mellins, E. D. Cathepsin G: roles in antigen presentation and beyond. Molecular immunology 47, 658–665 (2010).

76. Soman, A. & Asha Nair, S. Unfolding the cascade of SERPINA3: Inflammation to cancer. Biochimica et biophysica acta. Reviews on cancer 1877, (2022).

77. Wang, L., Yu, X., Zhou, J. & Su, C. Extracellular Vesicles for Drug Delivery in Cancer Treatment. Biological procedures online 25, (2023).

78. Vader, P., Breakefield, X. O. & Wood, M. J. A. Extracellular vesicles: emerging targets for cancer therapy. Trends in molecular medicine 20, 385–393 (2014).

